# A U1–U3 snRNA–snoRNA interaction couples SF3B1 mutation to chromatin-state rewiring and genome instability

**DOI:** 10.64898/2026.06.06.722969

**Authors:** Peng Xia, Han Li, Yiyi Ji, Cheng-wei Ju, Yiqian Pan, Jing Mo, Xuanhao Zhu, Lijie Zhao, Ruitu Lv, Erica Niewold, Mike Fernandez, Yuxi Ai, Jiangbo Wei, Robert K Bradley, Lili Wang, Omar Abdel-Wahab, Bei Liu, Chuan He

**Affiliations:** Department of Chemistry, The University of Chicago, Chicago, IL 60637, USA; Department of Biochemistry and Molecular Biology, The University of Chicago, Chicago, IL 60637, USA; Howard Hughes Medical Institute, The University of Chicago, Chicago, IL 60637, USA; Department of Systems Biology, Beckman Research Institute of the City of Hope, Monrovia, CA, USA; Irell Graduate School of Biological Sciences, Beckman Research Institute of the City of Hope, Duarte, CA, USA; Molecular Pharmacology Program, Sloan Kettering Institute, Memorial Sloan Kettering Cancer, New York, NY 10065, USA; Department of Chemistry, National University of Singapore, Singapore 119077, Singapore; Computational Biology Program, Public Health Sciences Division, Fred Hutchinson Cancer Research Center, Seattle, WA, USA; Herbert Irving Comprehensive Cancer Center, Columbia University, New York, NY 10032, USA

## Abstract

Mutations in spliceosome factors such as SF3B1 are recurrent across human diseases, including myelodysplastic syndromes and leukemia^1–4^, yet splicing defects alone do not fully explain the widespread chromatin alterations and genome instability in mutant cells^5^. Here, by comprehensively mapping snRNA-directed RNA–RNA interactions, we identify two previously unrecognized interaction motifs in U1 snRNA beyond canonical 5′ splice-site pairing^6,7^. These motifs enable U1 RNA to i) bind intronic and other chromatin-associated RNA (caRNA) regions outside of splice sites, and ii) base pair specifically with snoRNA. We uncover a U1–U3 snRNA–snoRNA interaction that recruits the H3K36 methyltransferase SETD2 to caRNA, promoting gene-body H3K36me3 and antagonizing H3K27me3 to modulate chromatin accessibility. The snRNA–snoRNA interface is essential for this previously unrecognized layer of chromatin and transcriptional regulation mediated through SETD2. SF3B1 mutation enhances U1–U3 binding and increases the association of the U1–U3 complex with caRNA, driving chromatin-accessibility remolding, R-loop formation, DNA damage, and copy-number abnormalities that promote tumorigenesis. A U1-specific 2′-O-methoxyethyl antisense oligonucleotide that selectively blocks U1–U3 pairing suppresses these genomic abnormalities, reduces leukemic infiltration, and prolongs survival in xenograft and patient-derived models, establishing pathological snRNA–snoRNA rewiring as a critical driver of SF3B1-mutant leukemogenesis.

## Main

Small nuclear RNAs (snRNAs) and small nucleolar RNAs (snoRNAs) are abundant noncoding RNAs that function as core cofactors for RNA processing. snRNAs are best known as integral components of the spliceosome, where they mediate recognition and catalysis at exon–intron junctions during pre-mRNA splicing^8^. snoRNAs, in contrast, predominantly localize to the nucleolus, where they mostly guide chemical modification of ribosomal RNAs and other nuclear RNAs^9,10^. Beyond these canonical processing roles, recent work has revealed that snRNAs and snoRNAs can influence transcriptional and chromatin regulation^11–13^. U1 snRNP (small nuclear ribonucleoprotein) exemplifies this expanded functionality: in addition to recognizing 5′ splice sites in pre-mRNA splicing, U1 can assemble with cleavage and polyadenylation (CPA) factors into a distinct U1–CPA factor complex (U1-CPAFs) that mediates long-range “telescripting” to restrain intronic polyadenylation signal (PAS) usage during transcription, thereby preserving transcript length and transcriptome integrity^7,14,15^. U1 snRNP has also been implicated in regulating the chromatin retention of noncoding RNAs^11^. In all of these settings U1 function depends on base pairing between the U1 snRNA and complementary sequences in its RNA targets.

Despite these advances, the binding sites and interaction networks of snRNAs have not been systematically mapped at a genome-wide scale. This lack of comprehensive maps creates a critical gap: without knowing where and how these RNAs engage potential target RNAs across the genome, it remains difficult to explain their non-canonical roles and potential involvements in human diseases.

Spliceosome factor mutations in human cancer illustrate this problem. Such mutations are particularly prevalent in leukemia^1,16^; in chronic lymphocytic leukemia (CLL), their frequency is estimated at 5–20%, where they are strongly associated with rapid disease progression and poor prognosis^5,17,18^. Among these, SF3B1 is a paradigmatic example^19,20^. SF3B1 is a core component of U2 snRNP that stabilizes U2 engagement at intronic branch site sequences during early spliceosome assembly, in coordination with U1 snRNP recognition of the 5′ splice site^4,17,21^. Although recurrent SF3B1 hotspot mutations such as K700E (SF3B1^K700E^) promotes the aberrant use of cryptic 3′ splice sites, global pre-mRNA splicing patterns are modestly affected^21^, suggesting additional pathogenic consequences of SF3B1 mutations beyond splicing defects.

An increasing body of evidence indicates that spliceosomal mutations also interfere with transcriptional dynamics and genome integrity^22^, leading to aberrant accumulation of RNA–DNA hybrids (R-loops) and subsequent DNA damage^23^. Although these mutations elicit well-defined splicing changes^21,24^, these splicing defects alone may not account for the observed extensive epigenetic remodeling in mutant cells. We hypothesized that spliceosome mutations may reshape chromatin organization through noncanonical molecular pathways. To explore this possibility, we investigated the potential role of spliceosomal RNAs in chromatin regulation. Here, we report a comprehensive map of snRNA‒target RNA interactions in mammalian cell lines. We discovered two previously unrecognized sequence motifs in U1 snRNA that mediate i) stable duplex formation with U3 snoRNA and ii) sequence-specific binding to chromatin-associated RNAs within gene bodies. This caRNA–U1–U3 RNA scaffold recruits the H3K36 methyltransferase SETD2 to modulate chromatin accessibility. This pathway is pathologically rewired in SF3B1-mutant leukemia. In SF3B1^K700E^ mutant cells we observed aberrant accumulation of U1–U3 complexes on chromatin, excessive SETD2 recruitment, pervasive R-loop formation, DNA damage, and copy-number alterations. We further show that a U1-specific, steric-blocking antisense oligonucleotide (ASO) that disrupts U1–U3 pairing attenuates these chromatin and genome-instability phenotypes, reduces leukaemic burden, and prolongs survival in xenograft and patient-derived xenograft models.

### Mapping snRNA–RNA interaction networks by snRNA-KARR-seq

To comprehensively map snRNA binding sites, we established kethoxal-assisted, snRNA-mediated RNA–RNA interactome sequencing (snRNA-KARR-seq), adapted from our previously developed KARR-seq method^25,26^. This approach enriches crosslinked snRNA complexes using snRNA-targeted antisense oligonucleotides (snRNA-ASOs), followed by conversion into chimeric cDNA libraries, thereby enabling high-throughput mapping of RNA–RNA interactions anchored on specific snRNAs (**Extended Data Fig. 1a**). Applying this method to multiple cell lines (HEK293, HepG2, K562 and HG3), we successfully generated high-quality libraries with high reproducibility (**Extended Data Fig. 1b**). Sequencing results revealed significant enrichment of snRNA reads and increased proportions of chimeric reads of snRNAs when using ASO for snRNA enrichment, confirming efficient and specific capture of snRNA-associated RNA interactions (**Extended Data Fig. 1c**,**d**).

As a benchmark, all captured snRNAs showed clear peaks at 5′ and 3′ splice sites (**Fig. 1a**), consistent with their canonical roles in pre-mRNA splicing. The previously characterized snRNA-target interactions, including U1-MALAT1 and U1-SCARNA16 interactions required for U1 chemical modifications (pseudouridine at position 5), were also successfully captured (**Extended Data Fig. 2a**). In addition to its canonical role in pre-mRNA splicing, U1 snRNP suppresses PCPA at cryptic intronic PAS^7,14^. Consistent with this, snRNA-KARR-seq data revealed U1 occupancy in the vicinity of PAS (**Extended Data Fig. 2b**,**c**). Therefore, our datasets provided a framework for systematically interrogating snRNA-directed RNA–RNA networks beyond canonical splice-site pairing.

**Fig 1.**
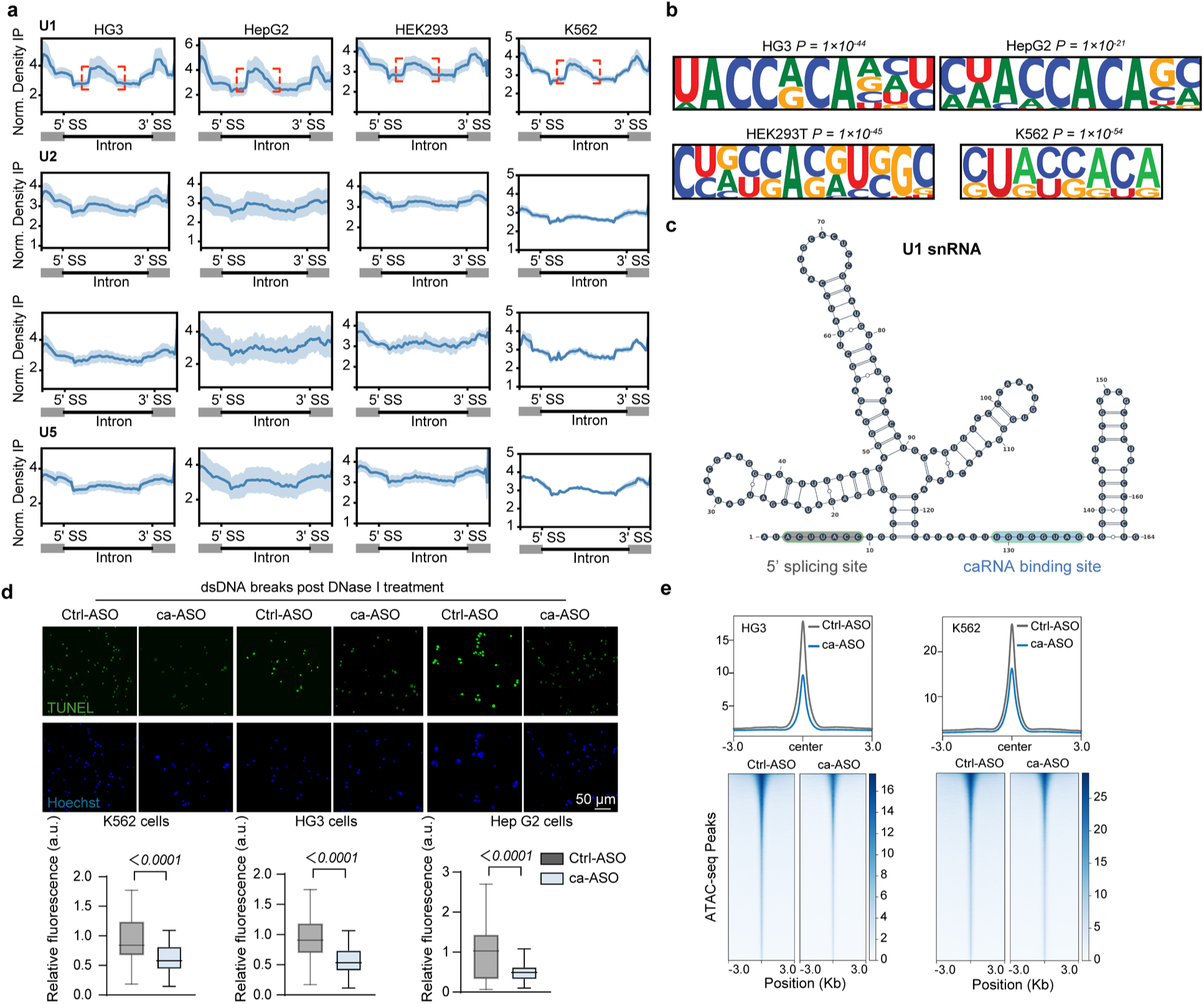
Chromatin accessibility and U1 snRNA binding motifs. **a**, Metagene profiles of snRNA binding sites across intronic regions and in proximity to splicing junctions, including the 5ʹ splice site (5ʹSS) and 3ʹ splice site (3ʹSS) across K562, HG3, HepG2, and HEK293 cell lines. **b**, Sequence motifs enriched in intronic regions bound by U1 snRNA in K562, HG3, HepG2, and HEK293 cell lines. **c**, Secondary structure of U1 snRNA, with the 5ʹ splice junction binding motif highlighted in grey and the caRNA interaction region in blue. **d**, Representative DNase I and TUNEL staining in Ctrl-ASO– and ca-ASO–treated K562, HG3, and HepG2 cells (left), with quantification across six fields of view (right; scale bar, 50 μm). **e**, Average ATAC-seq profiles and heat maps of chromatin accessibility in Ctrl-ASO– and ca-ASO–treated K562 and HG3 cells. For **d**, the box plots show the median (centre line), upper and lower quartiles (box limits) and 1–99% (whiskers). *P* value was determined using an unpaired two-tailed *t* test (**d**). TUNEL intensity was quantified by ImageJ.

Notably, a careful examination of the binding landscape of different snRNAs revealed to us that U1 displays extensive enrichment across intronic regions beyond splice sites, a feature not observed for other snRNAs across different cell lines (**Fig. 1a** and **Extended Data Fig. 2d**,**e**). Given that cellular snRNA abundances do not match their equimolar use in the spliceosome, and that U1 far exceeds other snRNAs in higher eukaryotes^14^, this observation indicated additional functions, such as telescripting, beyond its splicing role. Our analysis of U1 interactions with chromatin-associated RNAs (caRNA) identified a previously unrecognized, conserved intronic binding motif, “UACCACA,” which is highly preserved across multiple human cell lines (**Fig. 1b**). Importantly, this motif is distinct from the 5′ sequences recognized by U1 during classical splicing and proposed for transcriptional telescripting^7,14^ (**Fig. 1c**), suggesting that U1 forms a widespread intronic RNA interaction network and may exert additional functions on chromatin.

To further probe this possibility, we designed a 2′-O-methoxyethyl (2ʹ-MOE)–modified, steric-blocking ASO (ca-ASO) that targets the caRNA-binding sequence of U1 snRNA to disrupt the interaction between U1 and caRNA intronic sites. Flow cytometry confirmed efficient ca-ASO transfection, and qRT–PCR showed that ca-ASO treatment did not alter U1 expression level (**Extended Data Fig. 3a**,**b**). Previous studies showed that a U1 ASO targeting the 5′ splice-site recognition region (5′ splice-ASO) disrupts both U1-dependent splicing and suppression of PCPA^14^. To test whether ca-ASO has similar effects, we first examined *in vitro* splicing using ca-ASO with 5′ splice-ASO as a control. 5′ splice-ASO markedly reduced formation of the spliced Ad2ΔIVS product, whereas ca-ASO had no detectable impact on splicing (**Extended Data Fig. 3c**). In K562 cells, transfection with a 5′ splice-ASO induced widespread changes in differential splicing events, whereas the ca-ASO elicited only a limited number of alterations (**Extended Data Fig. 3d**). We further examined intronic polyadenylation (IPA) site usage in K562 cells and found that both the 5′ splice-ASO and the ca-ASO altered IPA site utilization (**Extended Data Fig. 3e**). This indicates that U1 binding to these caRNA intronic sites contributes to PCPA suppression in addition to its binding to cryptic 5′ splice sites, confirming a previous speculation that U1 snRNP may base pair to sequences other than the 5′ splice site to suppress PCPA^14^.

We next assessed whether this newly identified U1–intronic caRNA engagement has additional functional consequences. In K562, HG3, and HepG2 cells, ca-ASO treatment led to a marked reduction in chromatin accessibility, as demonstrated by DNase sensitivity and TUNEL assays (**Fig. 1d**). ATAC-seq confirmed a global shift toward chromatin closure, with the most pronounced effects occurring at caRNA regions corresponding to U1-binding sites identified by snRNA-KARR-seq (**Fig. 1e**). Together, these findings indicate that U1 snRNA helps maintain chromatin accessibility through binding to the UACCACA motif within caRNA, primarily in intronic regions. We next sought to delineate the molecular pathway underlying this activity and to determine its functional relevance.

### Identification of a stable U1–U3 snRNA-snoRNA duplex conserved across cell types

We subsequently examined the classes of RNAs bound by snRNAs. Interestingly, besides snRNA‒caRNA interactions, small nucleolar RNA (snoRNA) is another major target of snRNAs (**Fig. 2a**). snoRNAs are traditionally known to function as guides for site-specific ribosomal RNA modifications in order to facilitate accurate rRNA processing and ribosome assembly^9,10^. The identification of widespread and stable snRNA–snoRNA interactions (**Fig. 2a** and **Extended Data Fig. 4a**) may suggest a role for snoRNAs beyond their involvement in ribosome biogenesis.

**Fig 2.**
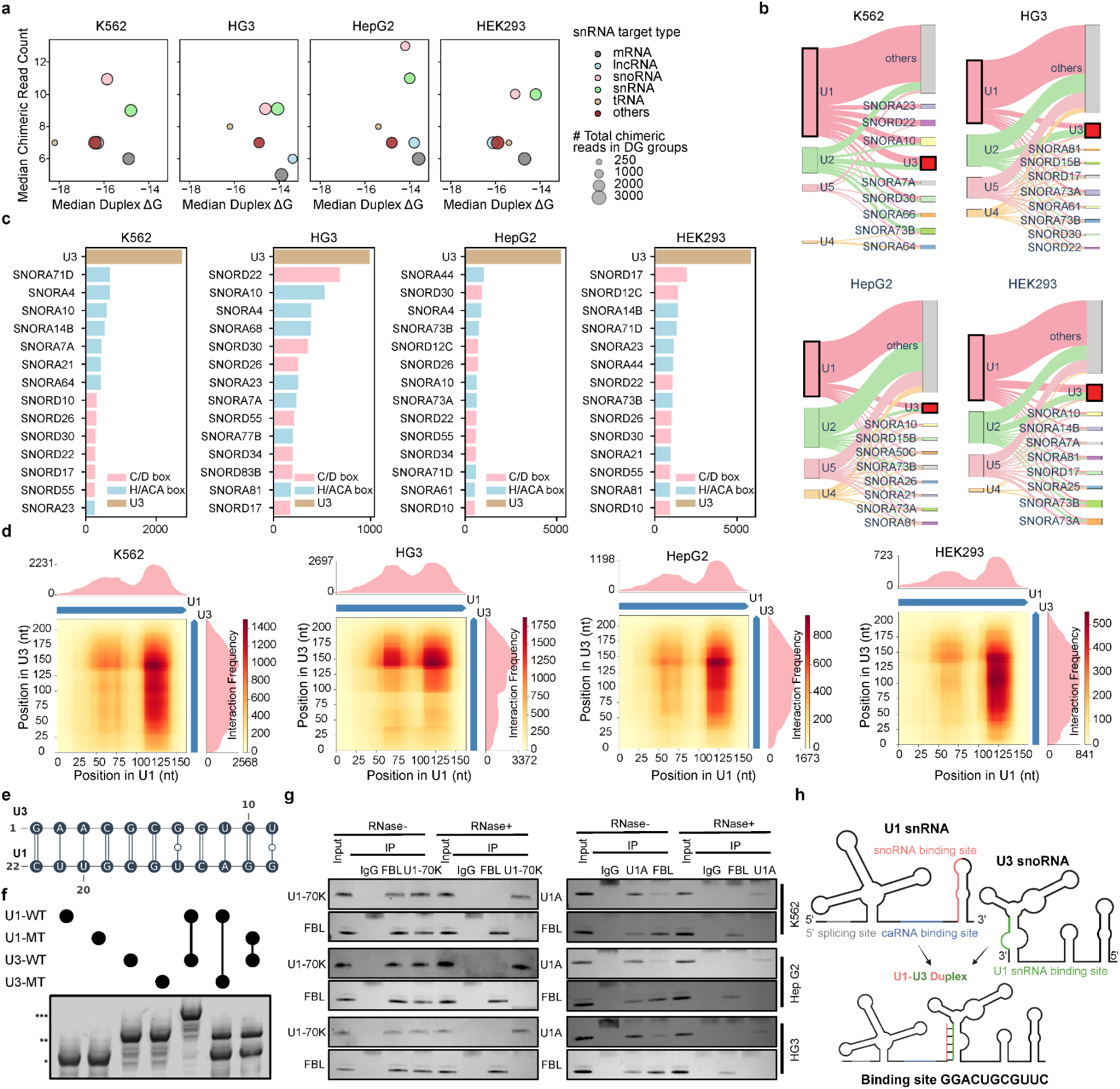
Identification and validation of a stable U1–U3 duplex. **a**, Quantification of chimeric reads and corresponding median duplex free energy (ΔG, kcal/mol) across distinct categories of snRNA target interactions in K562, HG3, HepG2, and HEK293 cells. **b**, Distribution patterns of snRNA-snoRNA interactions across K562, HG3, HepG2, and HEK293 cells. **c**, Chimeric read abundance for individual snoRNAs targeted by U1 snRNA across K562, HG3, HepG2, and HEK293 cells. **d**, Heatmap visualizing interaction frequencies between U1 and U3 snoRNAs across K562, HG3, HepG2, and HEK293 cells. **e**, Schematic representation of the U1–U3 base-pairing interface. **f**, Native gel electrophoresis showing formation of the U1–U3 duplex. In vitro–transcribed U1 and U3 RNAs, together with binding-site mutants, were annealed and analyzed by gel shift. The shifted band indicates duplex formation. *, U1; **, U3; ***, U1–U3 duplex (*n* = 3 biological replicates). **g**, Endogenous co-immunoprecipitation following UV crosslinking in K562, HG3, and HepG2 cells, performed with or without RNase treatment, with Western blot detection of interactions between U1-associated proteins (U1A and U1-70K) and the U3-associated protein FBL (*n* = 3 biological replicates). **h**, Schematic of the U1–U3 RNA duplex and the corresponding binding-site sequences.

Notably, these interactions are dominated by U1 snRNA, which formed the majority of detectable complexes with snoRNAs (**Fig. 2b**). Quantitative analysis further revealed that among all U1-associated snoRNAs, U3 produced the highest frequency of chimeric reads and strong duplex stability, greatly exceeding other candidates (**Fig. 2c**,**d** and **Extended Data Fig. 4b**).

Sequence mapping revealed a highly consistent and recurrent interaction interface, where nucleotides at the U1 3′ end (GGACUGCGUUC) form stable base pairing with complementary sequences (GAACGCGGUCU) in the 3′ domain of U3 (**Fig. 2d**,**e** and **Extended Data Fig. 4c–e**). This distinct binding pattern indicates the formation of a structured RNA duplex, rather than transient contacts. Importantly, the U1–U3 interaction was consistently captured across all tested cell lines, including K562, HEK293, HG3, and HepG2, underscoring its generality and conservation across cellular contexts (**Fig. 2d** and **Extended Data Fig. 4b**,**c**). Structure predictions guided by snRNA-KARR-seq chimeric reads revealed that the lowest free-energy U1-U3 duplexes consistently mapped to the same base-pairing regions across multiple cell types (**Extended Data Fig. 4c**), suggesting a conserved and potentially functionally relevant structural interface.

We next validated the physical interaction between U1 and U3 using orthogonal approaches. *In vitro* RNA duplex assays were performed with wild-type (WT) and mutant variants of U1 and U3, in which the predicted base-pairing regions were disrupted. When U1-WT and U3-WT transcripts were annealed, a distinct, slower-migrating complex appeared, consistent with duplex formation. In contrast, disruption of the complementary sequences in either U1 or U3 abolished complex formation, indicating strict dependence on sequence-specific base pairing (**Fig. 2e**,**f**). We next used UV crosslinking followed by endogenous co-immunoprecipitation to assess this association. Antibodies against the U1 snRNP components U1A and U1-70K co-precipitated fibrillarin (FBL), a core U3 snoRNP protein^9,27^. Reciprocally, FBL immunoprecipitation co-precipitated well-known U1 binding proteins U1A and U1-70K^28,29^. Importantly, this co-precipitation was abolished by RNase treatment, indicating that the observed co-precipitation relies on direct RNA–RNA base pairing rather than indirect protein–protein associations (**Fig. 2g**). Consistent with these results, RNA immunoprecipitation (RIP)–qPCR performed in the chromatin fraction using U1A/U1-70K and FBL, respectively, further demonstrated significant reciprocal enrichment, with U1-associated proteins pulling down U3 snoRNA and FBL enriching for U1 snRNA (**Extended Data Fig. 4f**). Collectively, these results establish a specific and stable U1–U3 snRNA–snoRNA duplex formation, which is also present in the chromatin fraction (**Fig. 2h**).

### The U1–U3 duplex remodels chromatin by promoting H3K36me3 deposition

To probe the functional role of the U1–U3 RNA duplex, we employed steric-blocking 2ʹ-MOE–modified ASOs targeting either U1 snRNA or U3 snoRNA to disrupt the U1–U3 pairing interface. Transfection efficiency was verified by flow cytometry (**Extended Data Fig. 5a**). qRT-PCR confirmed that overall U1 and U3 RNA levels remained stable following ASO treatment, indicating that the downstream effects reflect disruption of U1–U3 RNA–RNA interaction rather than RNA degradation (**Extended Data Fig. 5b**). DNase I sensitivity assay combined with TUNEL staining revealed a widespread reduction in open chromatin signals in both U1-ASO- and U3-ASO-treated cells (**Fig. 3a**). ATAC-seq analysis reinforced this observation, revealing a marked loss of chromatin accessibility following U1-ASO treatment (**Fig. 3b** and **Extended Data Fig. 5c**). Notably, regions exhibiting the strongest accessibility reductions tended to lie in close proximity to high-affinity caRNA U1-binding sites identified by snRNA-KARR-seq (**Fig. 3c** and **Extended Data Fig. 5d**). The magnitude of ATAC signal loss showed a clear distance-dependent pattern: loci within only a few kilobases of a U1-binding site displayed the most pronounced decreases, whereas more distal regions were progressively less affected (**Fig. 3c** and **Extended Data Fig. 5d**). The highly concordant peak-loss profiles induced by U1-ASO and U3-ASO are inconsistent with canonical, independent functions of U1 in pre-mRNA splicing and IPA site utilization, or U3 in rRNA processing, as we observed neither the widespread splicing perturbations nor the rRNA expression changes (**Extended Data Fig. 5e–j**). Together, these findings indicate that the U1–U3 complex contributes directly to maintaining an open chromatin state through a noncanonical function.

**Fig 3.**
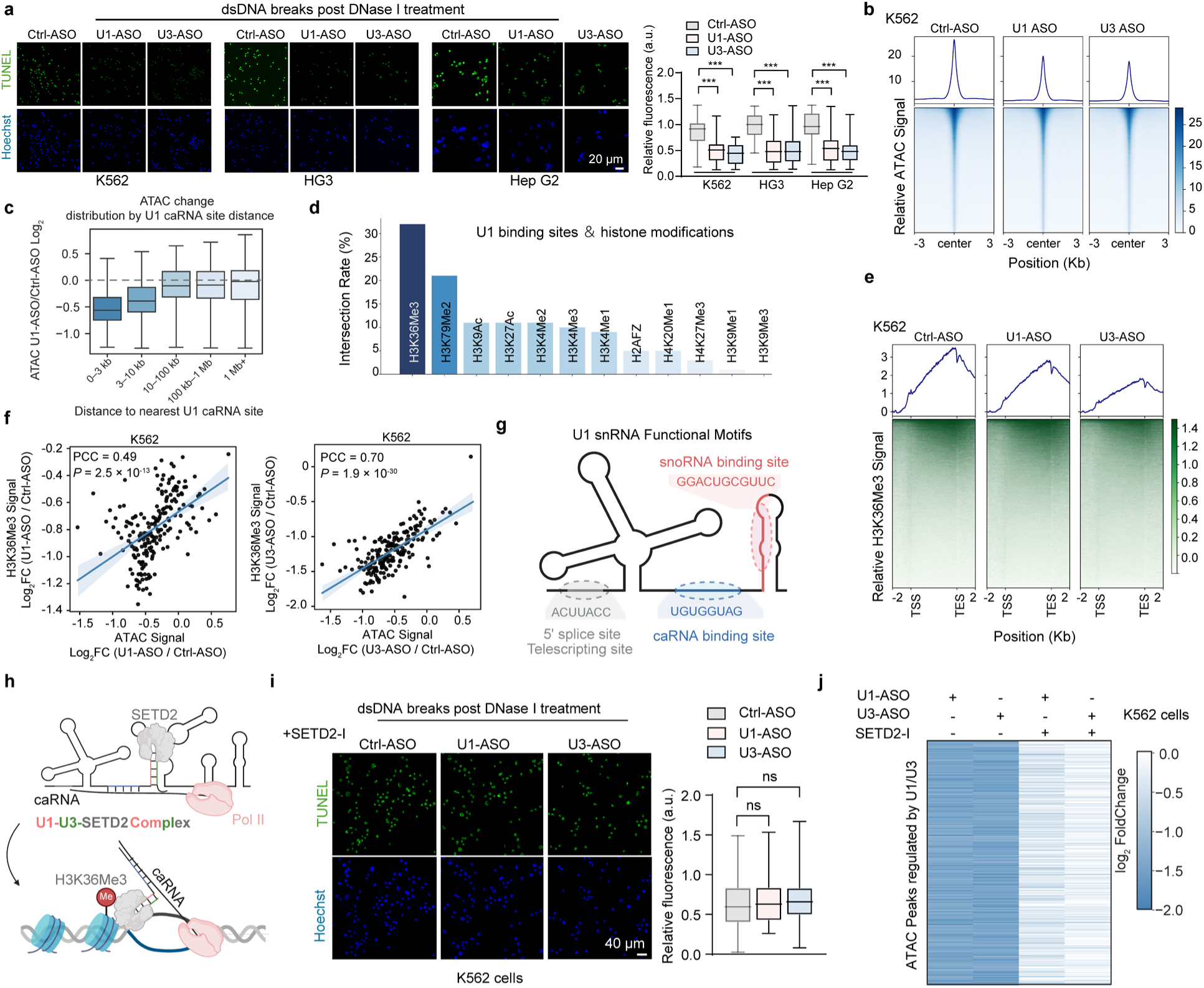
The U1–U3 duplex recruits SETD2 to promote H3K36me3 deposition and modulate chromatin accessibility. **a**, Representative DNase I and TUNEL staining in K562, HG3, and HepG2 cells showing changes in chromatin accessibility following disruption of U1–U3 binding by U1-ASO or U3-ASO (left), with quantification across six fields of view (right; scale bar, 20 μm). **b**, Average ATAC-seq profiles and heat maps of chromatin accessibility in K562 cells following U1-ASO or U3-ASO treatment. **c**, ATAC-seq signal changes showed a pronounced spatial gradient, with the largest reductions occurring within a few kilobases of U1–caRNA binding sites and effects diminishing toward zero at more distal regions in K562 cells. **d**, Overlap between U1-binding caRNA regions identified by snRNA-KARR-seq and histone modification signals from ENCODE ChIP-seq datasets in K562 cells. **e**, CUT&Tag signal changes in K562 cells showing changes in H3K36me3 peaks after U1-ASO or U3-ASO treatment. **f**, Correlation analysis in K562 cells between H3K36me3 peaks and ATAC peak alterations following U1-ASO or U3-ASO treatment. **g**, Schematic illustrating distinct sequence elements on U1 snRNA that mediate binding to snoRNAs sites, 5′ splice sites, and caRNA binding site. **h**, Schematic illustrating how the U1–U3 duplex recruits SETD2 through RNA binding to deposit H3K36me3 and maintain chromatin accessibility. **i**, Representative DNase I and TUNEL staining images showing chromatin alterations induced by U1-ASO or U3-ASO treatment upon SETD2 inhibition in K562 cells (left; scale bar, 40 μm). Quantification from six independent fields is shown on the right. **j**, Heat map of differential ATAC peaks in K562 cells after U1-ASO or U3-ASO treatment with or without SETD2 inhibition. *n* = 3 biological replicates. For **a**, **c**, the box plots show the median (centre line), upper and lower quartiles (box limits) and 1–99% (whiskers). *P* value was determined using an unpaired two-tailed *t* test (**a**), and Pearson’s correlation coefficient (PCC) was used to assess correlation (**f**). ns indicates no statistically significant difference. TUNEL intensity was quantified by ImageJ.

We next assessed whether these chromatin changes were accompanied by alterations in histone modifications. We noticed that U1 binding sites on caRNA showed substantial overlap with regions enriched for H3K36me3 (**Fig. 3d**). Western blotting showed that disrupting the U1–U3 duplex selectively reduced H3K36me3 levels (**Extended Data Fig. 6a**). CUT&Tag profiling of H3K36me3 in cells treated with control ASO versus U1-ASO or U3-ASO revealed a striking redistribution of this histone mark. In control cells, H3K36me3 was prominently enriched across intronic regions of actively transcribed genes. However, disrupting the U1–U3 pairing led to a widespread loss of H3K36me3 peaks across the genome (**Fig. 3e** and **Extended Data Fig. 6b**). Notably, the regions showing the strongest reductions in H3K36me3 closely mirror those exhibiting decreased chromatin accessibility in the ATAC-seq analysis following U1–U3 disruption (**Fig. 3f** and **Extended Data Fig. 6c**). This convergence supports that chromatin accessibility and H3K36me3 deposition lie on the same regulatory axis, both responding in parallel to disruption of the U1–U3 interaction.

Further stratification by distance to the nearest U1–caRNA overlap site revealed a clear positional effect on H3K36me3 loss. Loci located within only a few kilobases of a caRNA U1-binding site showed the strongest reductions, whereas regions progressively farther away displayed attenuated changes that approached zero (**Extended Data Fig. 6d**). This distance-dependent pattern indicates that the impact of U1–caRNA disruption on H3K36me3 is locally concentrated and diminishes with genomic separation. Notably, direct inhibition of U1–caRNA binding produced a similar distance-dependent decrease in H3K36me3, indicating a localized role for U1–caRNA engagement in positioning and maintaining H3K36me3 deposition (**Extended Data Fig. 6e–g**).

H3K36me3 is a critical determinant of chromatin homeostasis, and both its loss and its ectopic accumulation can perturb chromatin states, underscoring the need for tight control over its abundance and genomic distribution^30–32^. One established function of H3K36me3 is to facilitate the removal of elongation-associated acetylation and thereby suppress aberrant intragenic promoter activity^33,34^. In parallel, nucleosomal H3K36me3 directly antagonizes PRC2 catalysis, disfavoring H3K27me3 deposition on transcription-associated chromatin^35–37^. The functional relationship between H3K36me3 and chromatin accessibility is likely to be highly context- and locus-dependent, and the extent to which H3K36me3 promotes or constrains chromatin openness in different genomic contexts remains to be further investigated.

Altogether, our results so far revealed a new role of U1 snRNA in chromatin regulation. It apparently contains three distinct sequences involved in RNA–RNA interactions (**Fig. 3g**). The canonical sequence of ACUUACC at 5′ splice-site to recognize splicing sites on pre-mRNA, and recognize pre-mRNA in its telescripting role, and the two sequences discovered in this study: UGUGGUAG at mid-region of U1 for binding to caRNA (primarily introns) and GGACUGCGUUC at 3′-terminal of U1 for U3 specific binding (**Fig. 3g**). Our data further identify the U1–U3 RNA duplex as a bona fide regulatory RNA–RNA module that maintains chromatin accessibility across gene bodies genome-wide, as selective disruption of this RNA–RNA pairing reduces accessibility at U1 caRNA-binding sites. This mechanism represents a previously unrecognized RNA-based pathway of chromatin regulation, unique from prevailing models that emphasize protein–DNA interactions in epigenetic control.

### The U1–U3 duplex directly recruits SETD2 to mediate RNA-guided histone modification

H3K36me3 is deposited co-transcriptionally by the histone methyltransferase SETD2, the sole mammalian enzyme installing this modification^38^. The strong concordance between U1–U3 disruption and loss of H3K36me3 prompted us to test whether SETD2 is directly involved. We hypothesized that the U1–U3 duplex docked on caRNA may recruit SETD2 at transcribed chromatin, thereby facilitating H3K36me3 deposition (**Fig. 3h**). Quantitative fluorescence polarization (FP) and surface plasmon resonance (SPR) assays confirmed this high-affinity interaction, with an apparent dissociation constant (KD) of ∼40 nM for the U1–U3 duplex, compared with ∼400 nM for U1 alone and negligible binding to U3 (**Extended Data Fig. 7a**). These results indicate a strong and specific binding of SETD2 for the base-paired U1–U3 duplex. To define the SETD2 interaction domain, we generated a series of truncation constructs spanning distinct structural regions of the protein. Consistent with our initial mapping, binding assays showed that fragments encompassing the C-terminal portion of SETD2 (amino acids 1401–2564) are sufficient to interact with the U1–U3 duplex (**Extended Data Fig. 7b**,**c**). Further dissection revealed that the AWS and SET domains are required for this interaction, whereas the WW domain is dispensable (**Extended Data Fig. 7d**). These results suggest that the U1–U3 duplex acts as an RNA scaffold that recruits SETD2 through its AWS and SET domains, thereby influencing chromatin accessibility.

We next asked whether the chromatin effects of U1–U3 disruption require SETD2 (**Extended Data Fig. 7e**). When SETD2 activity was blocked, treatment with U1- or U3-ASOs no longer produced any additional changes in chromatin state (**Fig. 3l** and **Extended Data Fig. 7f**). Consistent with this, ATAC-seq and CUT&Tag revealed that when SETD2 activity was blocked, U1- or U3-ASO treatment no longer produced the characteristic loss of H3K36me3 or ATAC-seq signal observed under normal conditions (**Fig. 3m** and **Extended Data Fig. 7g**,**h**). These results demonstrate that SETD2 is required for the chromatin-opening function of the U1–U3 duplex.

Together, these results support our model of the snRNA-guided chromatin activation: U1 snRNA and U3 snoRNA form a complementary RNA duplex that functions as a structural scaffold to directly recruit SETD2, thereby promoting local H3K36me3 deposition and antagonizing H3K27me3 to sustain an open chromatin state, with U1 snRNA also anchoring the U1–U3 complex on caRNA loci that can form U1–caRNA interactions (**Fig. 3h**).

### SF3B1 defects rewire the U1–U3 axis to drive chromatin abnormalities and genome instability

We next asked whether this phenomenon has functional relevance in a disease context. To this end, we performed snRNA-KARR-seq in disease models carrying spliceosomal mutations. We observed a pronounced elevation of U1–U3 chimeric read abundance in SF3B1-mutant cells (**Fig. 4a** and **Extended Data Fig. 8a**). Consistently, RIP–qPCR showed a significant increase in the association between U3 and U1 in SF3B1-mutant cell lines (**Fig. 4b**), suggesting that the U1–U3 duplex forms more readily in this setting. Notably, stem–loop 4 (SL4) of U1 snRNA is relatively exposed, and prior work has suggested that this region can engage SF3-associated components, with potential consequences for spliceosome assembly and snRNP dynamics^39^. We therefore reasoned that SF3B1 mutations might alter interactions between U1 and snoRNAs. In support of this idea, CLIP–seq of SF3B1 revealed an overall reduction in enrichment over U1 snRNA and its isoforms in SF3B1-mutant cells when compared to SF3B1 wild-type (**Extended Data Fig. 8b**). IGV inspection further indicated diminished binding signals of mutant SF3B1 over the 3′ region of U1 snRNA in mutant cells (**Extended Data Fig. 8c**), suggesting an increased accessibility of this U1 RNA region that could participate in snoRNA pairing. Concordantly, RIP–qPCR detected reduced association between mutant SF3B1 and U1 snRNA when compared with the wild-type SF3B1 (**Extended Data Fig. 8d**). Together, these findings propose a model in which SF3B1 mutation relaxes constraints on the 3′ end of U1 snRNA, thereby favoring formation of U1–U3 duplexes in chromatin-associated fractions.

**Fig 4.**
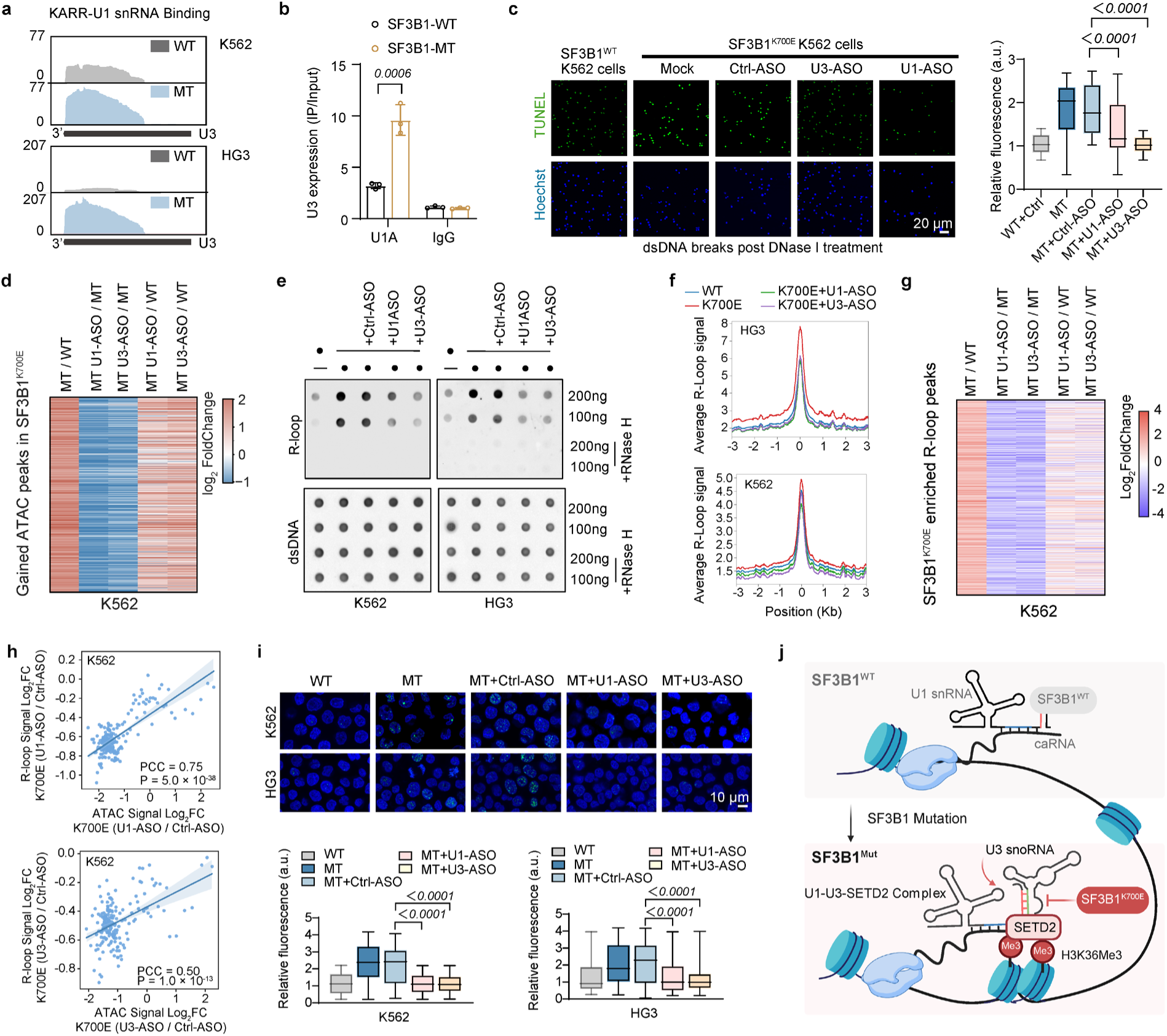
SF3B1 mutation amplifies U1–U3–dependent chromatin dysregulation and genome instability. **a**, Representative genome browser tracks showing U1 binding to U3 snoRNA loci in wild-type (SF3B1^K700K^) and mutant (SF3B1^K700E^) K562 and HG3 cell lines. **b**, RIP-qPCR showing enrichment of U3 snoRNA by U1A antibody in Flag-SF3B1^WT^ and Flag-SF3B1^K700E^ overexpression K562 cells. **c**, Representative DNase I and TUNEL staining in SF3B1^K700E^ K562 cells after U1-ASO or U3-ASO treatment (left), with quantification across six fields of view (right; scale bar, 20 μm). **d**, Heat map illustrating the spike-in-calibrated ATAC–seq signals across ATAC-seq peak regions that are more accessible in SF3B1^K700E^ K562 cells than in SF3B1^WT^ cells, and these signals change following U1-ASO or U3-ASO treatment. **e**, Dot blot analysis of R-loop levels in K562 and HG3 cells, showing elevated signals in SF3B1^K700E^ cells compared with SF3B1^K700K^ (WT) cells, and reduction upon U1-ASO or U3-ASO treatment. **f**, CUT&Tag profiling of R-loop–enriched regions show increased R-loop accumulation in SF3B1^K700E^ K562 or HG3 cells relative to SF3B1^K700K^ (WT) K562 or HG3 cells, and shows changes following U1-ASO or U3-ASO treatment. **g**, Heatmap showing spike-in–calibrated R-loop signals at R-loop peak regions that are increased in SF3B1^K700E^ K562 cells compared with SF3B1^K700K^ (WT) K562 cells, and their changes following U1-ASO or U3-ASO treatment. **h**, Correlation between peaks showing reduced ATAC-seq signal and reduced R-loop accumulation after ASO treatment in SF3B1^K700E^ K562 cells. **i**, Representative immunofluorescence staining for phosphorylated γ-H2AX and quantification of nuclear fluorescence intensity normalized to DAPI in K562 and HG3 cells (scale bar, 10 μm). Quantification from six independent fields is shown on the right. **j**, Schematic illustrating that SF3B1-mutant cells show aberrant accumulation of U1–U3 RNA duplexes, which recruit excess SETD2 to chromatin and drive elevated H3K36me3 deposition, resulting in abnormal chromatin opening. *n* = 3 biological per group. Data are mean ± s.d (**b**). For **c** and **i**, the box plots show the median (centre line), upper and lower quartiles (box limits) and 1–99% (whiskers). *P* value was determined using an unpaired two-tailed *t* test (**b** and **i**), and Pearson’s correlation coefficient (PCC) was used to assess correlation (**h**). TUNEL intensity was quantified by ImageJ.

SF3B1 is one of the most frequently mutated splicing factors in hematologic malignancies and is closely associated with reduced overall survival in CLL^3,5,19,40^. Given the increased abundance of the U1–U3 duplex in chromatin fractions from SF3B1-mutant cells, we next evaluated whether this aberrant accumulation is linked to malignant phenotypes associated with SF3B1 mutation. We first examined chromatin architecture in *SF3B1^K^*^700^*^E^* leukemic cells. DNase I digestion combined with TUNEL staining revealed that *SF3B1* mutant cells displayed globally enhanced chromatin accessibility (**Extended Data Fig. 8e**). ATAC-seq profiling further confirmed a genome-wide increase in open chromatin signals, with particularly pronounced accessibility in intron regions (**Extended Data Fig. 8f**,**g**). Western blotting revealed a pronounced increase in H3K36me3, accompanied by a reduction in H3K27me3, in SF3B1-mutant cells relative to SF3B1-WT cells (**Extended Data Fig. 8h**). This is consistent with previous ChIP–seq datasets showing that SF3B1 mutations lead to a global increase in intronic H3K36me3 levels^41^. Together, these results suggest that SF3B1^K700E^ not only perturbs splice-site choice, but is also associated with broader chromatin remodeling.

In addition, S9.6 immunofluorescence revealed widespread R-loop foci in SF3B1-mutant cells (**Extended Data Fig. 8i**), consistent with prior reports linking SF3B1 mutations to elevated R-loop formation^42^. CUT&Tag analysis incorporating spike-in normalization confirmed that SF3B1-mutant cells exhibited a markedly higher number of upregulated R-loop signals than downregulated ones (**Extended Data Fig. 8j**). Integrated analysis further showed that these newly formed R-loop peaks positively correlate with regions of increased chromatin accessibility detected by ATAC-seq in SF3B1-mutant cells (**Extended Data Fig. 8k**). The accumulation of R-loops provides a mechanistic link to genome instability, as R-loops impede replication fork progression and promote DNA breaks^42^. In line with this, SF3B1-mutant cells displayed significantly increased γH2AX foci intensity and frequency compared with SF3B1-WT cells, indicative of elevated DNA double-strand breaks (**Extended Data Fig. 8l**).

To assess whether aberrant U1–U3 duplex formation contributes to these phenotypes, we performed rescue experiments. Disruption of U1–U3 pairing with ASOs in SF3B1^K700E^ cells (K562 and HG3) largely restored chromatin accessibility to SF3B1-WT levels, as measured by ATAC-seq and DNase sensitivity (**Fig. 4c**,**d** and **Extended Data Fig. 9a–c**). Regions in which U1- or U3-ASO treatment reduced ATAC signals were strongly inversely correlated with regions that became more accessible in SF3B1-mutant cells relative to wild type, indicating that ASO-mediated disruption of the duplex effectively reverses the chromatin-opening effects of the mutation (**Extended Data Fig. 9d**). H3K36me3 patterns within intronic regions were similarly restored (**Extended Data Fig. 9e**). Moreover, S9.6 dot blot analysis combined with genome-wide R-loop profiling showed that disrupting the U1–U3 duplex almost completely eliminated the SF3B1 mutation–induced R-loop peaks, restoring R-loop levels to near–wild-type levels (**Fig. 4e–g** and **Extended Data Fig. 9f**). These results demonstrate that blocking the U1–U3 interaction effectively alleviates the aberrant R-loop accumulation driven by the SF3B1 mutation. Consistently, R-loop peaks reduced by ASO treatment are significantly inversely correlated with those abnormally enriched in SF3B1^K700E^ cells relative to wild type (**Extended Data Fig. 9g**).

To determine whether these R-loop changes reflect altered H3K36me3 deposition, we examined genome-wide relationships and found a strong positive correlation between ASO-reduced H3K36me3 and the decrease in R-loop signals (**Extended Data Fig. 9h**). Moreover, regions showing diminished R-loop accumulation also positively correlate with sites exhibiting reduced chromatin accessibility by ATAC-seq, indicating that alleviating aberrant U1–U3 deposition can mitigate both the chromatin-opening phenotype and R-loop formation (**Fig. 4h** and **Extended Data Fig. 9i**).

SF3B1 mutations are known to intensify R-loop accumulation and chronic DNA damage, thereby promoting genomic instability, driving sustained clonal evolution, increasing resistance to therapy, and ultimately leading to poorer clinical outcomes^20,43,44^. γH2AX immunofluorescence staining revealed a pronounced decrease in nuclear foci following U1–U3 disruption (**Fig. 4i**), indicating a substantial reduction in DNA damage. Importantly, disrupting the U1–U3 interaction significantly reduced SF3B1^K700E^-associated copy-number variants (CNVs), thereby mitigating genome instability in mutant cells (**Extended Data Fig. 9j**). Together, our findings reveal that SF3B1 mutation alters an RNA-guided chromatin modification pathway, rewiring the U1–U3–SETD2 axis to establish a chromatin state prone to genomic instability. Additionally, these results suggest a therapeutic opportunity: targeting U1–U3 interactions or their associated regulatory factors may selectively restore epigenetic homeostasis and reduce genomic instability in SF3B1-mutant malignancies (**Fig. 4j**).

### U1-ASO alleviates pathogenic phenotypes of SF3B1-mutant leukemia in vivo

The presence of SF3B1 mutations constitutes an independent risk factor for rapid disease progression^43,45^. Multiple strategies have been explored to correct aberrant splicing^4,6,24,46^. Genomic instability associated with these mutations is likely a critical driver of cancer initiation and progression. Our data indicate that dampening the SF3B1 mutation–induced accumulation of the U1–U3 complex on caRNA may effectively mitigate genome instability. Because U1-ASO selectively blocks U1–U3 pairing with minimal effects on splicing or PCPA (**Extended Data Fig. 5h–j**), we evaluated U1-ASO activity in SF3B1^K700E^ leukemia models. Two human leukemia xenograft models were established in highly immunodeficient NSG (NOD.Cg-*Prkdc ^scid^ Il2rg ^tm1Wjl^* /SzJ) mice by intravenous injection of SF3B1^K700E^ K562 or SF3B1^K700E^ HG3 cells. Once leukemic burden was confirmed, mice were treated once weekly via tail-vein injection with Ctrl-ASO (2′-MOE, 5 mg/kg) or U1-ASO (2′-MOE, 5 mg/kg; **Fig. 5a**). Across both models, U1-ASO treatment markedly reduced leukemic burden in SF3B1^K700E^ xenografts, as evidenced by substantially lower bioluminescence throughout disease progression, and significantly extended overall survival compared with Ctrl-ASO–treated mice (**Fig. 5b,c**). In SF3B1^K700E^ xenografts, treatment also markedly reduced splenomegaly (**Fig. 5d** and **Extended Data Fig. 10a**). Histological examination of the liver revealed fewer metastatic leukemic foci in U1-ASO–treated SF3B1^K700E^ xenograft mice (**Fig. 5e, Extended Data Fig. 10b**). Consistently, flow cytometry confirmed a marked reduction in human leukemia cell burden across the spleen, liver, bone marrow, and peripheral blood in SF3B1^K700E^ xenograft mice (**Fig. 5f** and **Extended Data Fig. 10c**), indicating effective control of tumor load and extramedullary infiltration.

**Fig. 5.**
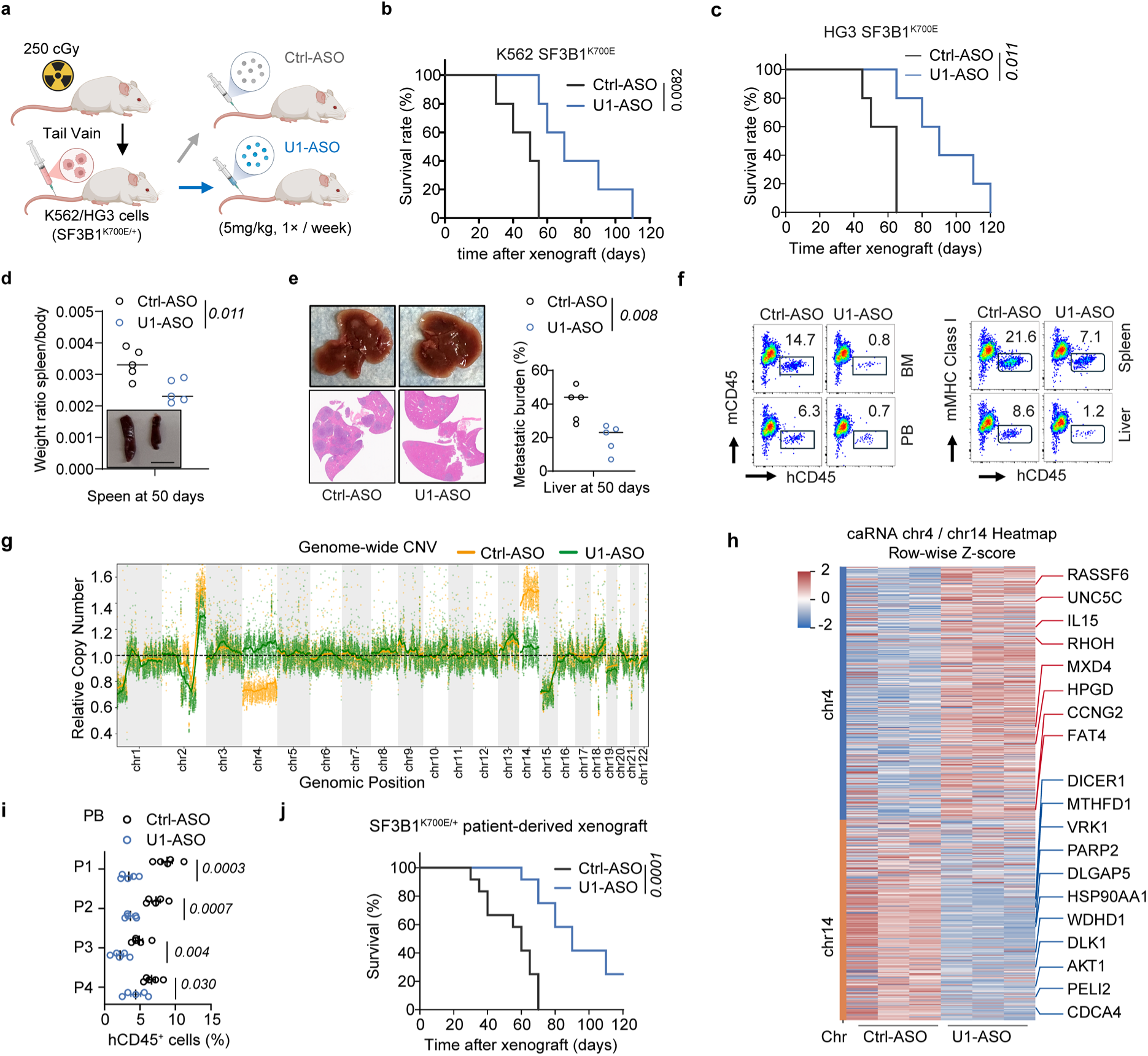
U1-ASO treatment suppresses leukemia progression and improves survival in SF3B1-mutant models. **a**, Schematic of xenograft experiments using luciferase-expressing K562 and HG3 cells with U1–U3 binding disrupted by U1-ASO. K562 cells or HG3 cells were injected intravenously into sublethally irradiated (250 cGy) NOD-scid IL2Rg^null^ (NSG) mice (*n* = 5 biological per group). **b**, Kaplan–Meier survival of mice bearing K562 xenografts. **c**, Kaplan–Meier survival of mice bearing HG3 xenografts. **d**, Representative spleen images and spleen weights from mice euthanized at day 50 after Ctrl-ASO or U1-ASO treatment in K562 xenograft model (*n* = 5 biological per group; scale bar, 1 cm). **e**, Number of hepatic metastatic foci in control-ASO– and U1-ASO–treated mice bearing K562 xenografts. **f**, Representative flow cytometry plots showing human leukemia K562 cells infiltration in peripheral blood, bone marrow, liver and speen (*n* = 5 biological per group). **g**, Whole-genome sequencing (WGS) analysis of copy-number variation (CNV) in K562 SF3B1^K700E^ cells recovered from mice 50 days after U1-ASO or Ctrl-ASO treatment. CNV profiles were derived from WGS read-depth data and normalized to Ctrl-ASO–treated K562 SF3B1^K700K^ WGS data to quantify genome-wide relative CNV. **h**, Heatmap of differential caRNA expression from chromosomes 4 and 14 in xenograft-derived K562 SF3B1^K700E^ cells, shown as row-wise z-scores. Representative upregulated and downregulated genes are highlighted with red and blue labels, respectively. **i**, Peripheral blood infiltration by four SF3B1-mutant human leukaemia samples in patient-derived xenograft (PDX) models at day 30 (*n* = 5 biological per group). **j**, Kaplan–Meier survival curves of PDX models carrying SF3B1^K700E^ mutation (*n* = 12 biological per group). *P* values computed by unpaired two-tailed t test (**d**, **e**, **i**) or Log-rank (Mantel–Cox) test (**c**, **c**, and **j**).

To further assess systemic safety, U1-ASO was administered at the same dose to healthy mice without leukemia engraftment (**Extended Data Fig. 10d**). This treatment did not result in abnormal changes in body weight or U1 expression, and histopathological examination of major organs revealed no evidence of tissue damage, inflammatory infiltration, or degenerative alterations (**Extended Data Fig. 10e–g**). Moreover, hematological parameters as well as liver and renal function indices remained comparable to those of untreated controls, with no hematologic or organ toxicities commonly associated with ASO treatment observed^47^ (**Extended Data Fig. 10h**). We therefore conclude that our U1-ASO treatment is well tolerated and exhibits a favorable safety profile in vivo.

To test whether U1-ASO mitigates genomic instability, we performed whole-genome sequencing (WGS) on sorted human leukemia cells from treated spleens. Compared with Ctrl-ASO group, U1-ASO substantially attenuated the SF3B1^K700E^-associated CNVs, most notably the gain of chromosome 14 and the loss of chromosome 4 (**Fig. 5g**). Blocking aberrant U1–U3 interactions reduced CNVs and clonal evolution, and this “deceleration” of genomic evolution may partially underlie the survival benefit conferred by U1-ASO and restrain leukemic expansion. To further support this conclusion, we performed caRNA-seq and mRNA profiling on tumour cells isolated from mice before and after U1-ASO treatment, focusing on CNV-affected chromosomes 4 and 14. In control tumours, these regions showed pronounced transcriptional rewiring, marked by induction of oncogenic genes and repression of tumour suppressors; U1-ASO treatment largely dampened these changes (**Fig. 5h** and **Extended Data Fig. 10i**). Together, these data indicate that disrupting aberrant U1–U3 interactions constrains the genomic and transcriptional consequences of SF3B1-driven instability.

We next evaluated U1-ASO efficacy in patient-derived xenograft (PDX) models. Four independent SF3B1^K700E^ leukemia samples, spanning distinct subtypes and genetic backgrounds, were engrafted into NSG mice. Across all SF3B1^K700E^ PDX cohorts, U1-ASO treatment significantly delayed disease progression and extended overall survival (**Fig. 5i**,**j**). Collectively, these data indicate that U1-ASO elicits consistent therapeutic activity in SF3B1-mutant leukemia models. By reducing leukemic burden, limiting extramedullary dissemination, and suppressing the selective advantage conferred by ongoing mutational accumulation, U1-ASO markedly prolongs host survival. These preclinical findings provide a strong rationale for targeting the U1–U3 RNA interaction as a therapeutic strategy for treating SF3B1-mutant leukemia.

## Discussion

snRNAs are central components of the spliceosome, directing intron recognition and excision during pre-mRNA splicing^47^, whereas snoRNAs guide site-specific rRNA modifications such as 2′-O-methylation and pseudouridylation to ensure ribosome maturation and function^26^. These two noncoding RNA classes have long been considered functionally independent. Our study uncovers an unexpected mode of crosstalk between U1 snRNA and U3 snoRNA and adds a new dimension to RNA-based chromatin regulation.

This discovery substantially broadens the functional repertoire of U1 snRNA beyond its canonical role in splicing and its known role in telescripting that prevents premature transcription termination and maintains transcriptional continuity^7,15^. U1 here uses two distinct new sequence motifs to form a stable duplex with U3 snoRNA and recognize chromatin-associated RNA, thereby anchoring the U1─U3 molecular scaffold in transcribing gene bodies to recruit the methyltransferase SETD2. We have recently discovered a class of snoRNAs that act as “*molecular glue*,” which connects the ribosome-nascent peptide-mRNA complex and the signal recognition particle through two distinct RNA─RNA interactions (mRNA─snoRNA─7SL RNA)^26^. Here, U1 snRNA serves as a functional hub that connects caRNA and U1 snRNA also through two distinct RNA-RNA interactions. This caRNA–U1–U3 module exemplifies a previously unrecognized RNA-guided mechanism that maintains proper gene-body H3K36me3 to sustain transcription.

This noncanonical role of U1 snRNA is consistent with emerging evidence that U1 can tether regulatory RNAs to chromatin^11^. Genome-wide RNA–chromatin mapping also uncovered widespread snRNAs and snoRNAs across chromatin^13^. Our identification of new sequence motifs embedded in U1 RNA that mediate extensive interactions with caRNA and U3 snoRNA supports a model that chimeric RNA–RNA structures function not only as recognition modules but also as sequence-dependent anchoring platforms for chromatin-modifying enzymes, thereby providing a new paradigm for RNA-mediated epigenetic regulation.

The significance of this pathway is highlighted in SF3B1-mutant leukemia, where spliceosome factor mutations are frequent drivers of malignancies^42^. SF3B1^K700E^ induces measurable splicing alterations, yet is paradoxically associated with prominent intronic chromatin abnormalities and genomic instability^21,40^. Our findings provide a mechanistic framework beyond splicing per se, implicating the U1–U3 chromatin pathway as a key downstream effector of the mutation. SF3B1 mutation perturbs snRNP dynamics, leading to aberrant accumulation of the U1–U3 duplex on chromatin. This causes aberrant SETD2 accumulation, disruption of the intronic H3K36me3–H3K27me3 balance, altered chromatin accessibility and pervasive R-loop formation—molecular features that directly connect the mutation to genome instability and tumor progression^23^. In this way, SF3B1 functions not only by altering branch-site selection but also by rewiring an RNA-guided chromatin modulation pathway. These insights extend the reach of spliceosome factor mutations into the epigenetic realm and illustrate how SF3B1 mutations exploit U1-centered RNA networks to erode genome integrity.

More broadly, these findings raise the possibility that snRNA-centred RNA–RNA interactions may constitute a general layer of regulation with additional, yet-to-be-defined functions. Our newly discovered U1‒intronic caRNA interaction should also impact telescripting (**Fig. 1d**), as suggested previously^14^, which requires future studies. Although we generated a comprehensive snRNA-centred RNA–RNA interaction map, we focused here on the functional consequences of U1 and its chromatin-associated interactions. It remains possible that additional snRNAs participate in analogous RNA–RNA networks and exert distinct regulatory functions that were not explored in this study. Furthermore, given the central role of the U1–U3 axis in chromatin homeostasis, perturbations beyond SF3B1 mutation may similarly disrupt this steady state and could be amenable to modulation with U1-ASO. However, we did not interrogate these alternative possibilities. Defining the scope of snRNA-centred interactions, their physiological and pathophysiological relevance, and their tractability as therapeutic targets will be important directions for future research^6,48^.

In summary, we identify an unprecedented role for a snRNA–snoRNA interaction in chromatin regulation. A functional RNA duplex between U1 and U3 acts as an RNA-based scaffold to recruit SETD2, coupling RNA-based sequence recognition to histone modification and chromatin architecture. Our findings position RNA–RNA interactions as critical regulators of epigenetic state, expand a paradigm historically dominated by protein–DNA interactions, and provide mechanistic insight into how SF3B1 mutations reprogram chromatin biology. Our in vivo data show that disruption of U1–U3 interactions with RNA-based agents effectively alleviates epigenomic and genomic abnormalities, reduces leukemic burden, and prolongs survival. Together, this work establishes noncoding RNA crosstalk as a key layer of chromatin regulation and provides a compelling rationale for targeting U1–U3 and related interactions as a future therapeutic strategy in SF3B1-mutant malignancies.

## Methods

### Animals

Female NSG (NOD.Cg-*Prkdc ^scid^ Il2rg ^tm1Wjl^* /SzJ) mice (6 weeks of age) were maintained under standard laboratory conditions with ad libitum access to food and water. Animals were randomly assigned to experimental groups, and all data collection and analyses were performed in a blinded manner by two independent laboratory members. SF3B1-mutant leukemia patient samples were obtained from the Omar Abdel-Wahab laboratory under written informed consent and with approval from the University of Chicago Institutional Review Board (IRB). All animal procedures were approved by the Institutional Animal Care and Use Committee (IACUC) of the University of Chicago (protocol no. 71847) and were conducted in strict accordance with institutional and national guidelines for the care and use of laboratory animals.

### Xenotransplantation of Leukemia Cells

For *in vivo* xenotransplantation assays, 1 × 10^6^ K562 or HG3 cells were intravenously injected via the tail vein into adult NSG mice (6–8 weeks old) that had been preconditioned with 250 cGy total-body irradiation. At day 50 (K562) or 60 (HG3) post-transplantation, peripheral blood (PB) was collected from the submandibular vein, and bone marrow (BM) cells were harvested from the femurs and tibias. Human CD45⁺ cell chimerism in each tissue was quantified using a BD FACSCelesta flow cytometer (BD Biosciences). Cells were stained with antibodies against human CD45 (hCD45), together with mouse CD45 (mCD45) or mouse MHC class I (mMHC-I), to determine the proportion of human leukemia cells in each tissue. Flow cytometry data were analyzed using FlowJo software (version 10.6.2).

### Cell Culture

K700K and K700E knock-in derivatives of K562 and HG3 cells were generated using previously described gene-editing procedures^40^. Wild-type HepG2 and HEK293T cell lines were obtained from the American Type Culture Collection (ATCC). Suspension cell lines, including K562 and HG3, were maintained in RPMI-1640 medium (Gibco, 61870036) supplemented with 10% fetal bovine serum (FBS; Gibco, 26140079) at 37 °C in a humidified atmosphere containing 5% CO_2_. Adherent HEK293T (ATCC, CRL-11268) and HepG2 (ATCC, HB-8065) cells were cultured in DMEM (Gibco, 11995). All media were supplemented with 10% (v/v) FBS and 1% penicillin– streptomycin (Gibco, 15140122). To ensure the absence of mycoplasma contamination, all cell lines used in this study were routinely screened with the LookOut Mycoplasma PCR Detection Kit (Sigma-Aldrich, MP0035).

### Synthesis of DBCO- and Biotin-Modified PAMAM G1 Dendrimers

To generate PAMAM G1 dendrimers simultaneously functionalized with DBCO and biotin, PAMAM G1 (1.53 μmol), DBCO-NHS (3.06 μmol), and biotin-NHS (1.53 μmol) were dissolved in 2 mL methanol, followed by the addition of 5 μL triethylamine to promote amide coupling. The reaction mixture was stirred gently at room temperature overnight to allow the activated esters to react with the primary amines on the dendrimer surface. Unreacted amines were subsequently capped by adding acetic anhydride (100 μL) and triethylamine (100 μL), and the mixture was stirred for an additional 24 h at room temperature. After completion of the reaction, 2 mL of water was added to quench the reaction, and the product was purified and concentrated by multiple centrifugation cycles at 5,000 g and 4 °C using a Microsep Advance ultrafiltration device equipped with a 1-kDa Omega membrane. The purified conjugates were quantified by measuring their characteristic absorbance at 295 nm, and the final DBCO content was determined using a calibration curve generated from serial dilutions of DBCO-NHS standards.

### ASO Design Targeting snRNA cDNA

ASOs targeting snRNA cDNAs were designed based on the corresponding snRNA sequences. snRNA sequences were retrieved from RNAcentral and converted to cDNA prior to probe design. ASO candidates were primarily generated using the Primer3 software; when suitable sequences could not be obtained, a complementary sequence of approximately 40 nucleotides from the central region of the snRNA cDNA was selected as an alternative. All ASOs were synthesized with a 5′ biotin modification by Integrated DNA Technologies (IDT). All ASO sequences used in this study are provided in **Supplementary Table 1**.

### snRNA-KARR-seq

snRNA-KARR-seq was performed as previously described with minor modifications^25^. Briefly, cells at approximately 80% confluence were crosslinked with 1% formaldehyde, quenched with glycine, and permeabilized in buffer containing N_3_-kethoxal. The permeabilized cells were then incubated with G1-DBCO-biotin at 37 °C. Following proteinase K digestion, RNA was isolated by phenol–chloroform extraction, ethanol precipitation, and DNase treatment, and subsequently subjected to fragmentation. Fragmented RNAs were enriched using Dynabeads MyOne Streptavidin C1 under high-salt conditions, dephosphorylated and rephosphorylated with T4 PNK, and ligated at the 3′ end using T4 RNA ligase. After elution and purification with the RNA Clean & Concentrator kit, reverse transcription was carried out using the SMARTer Stranded Total RNA-seq Kit v2. The resulting cDNAs were hybridized to biotinylated ASOs in a high-salt buffer using a stepwise temperature-decrease protocol, followed by a second streptavidin-based enrichment to isolate snoRNA-interacting sequences. Final libraries were constructed according to the manufacturer’s instructions and sequenced on an Illumina NovaSeq X platform using PE150 mode.

### DNase I–TUNEL Assay

The DNase I–TUNEL assay was performed using the DeadEnd Fluorometric TUNEL System (Promega, G3250) according to the manufacturer’s instructions. Cells were first fixed with paraformaldehyde and permeabilized with Triton X-100 to allow reagent access to the nuclei. For the positive control, cells were treated with DNase I (1 U μL⁻¹; Thermo Fisher Scientific, EN0521) at 37 °C for 5 min before the rTdT labeling reaction to induce DNA strand breaks. Following TUNEL labeling, fluorescence images were acquired using a Leica confocal microscope. All experiments were conducted in three independent biological replicates to ensure reproducibility.

### ATAC-seq

ATAC-seq was performed using the Hyperactive ATAC-Seq Library Prep Kit for Illumina (TD711) following the manufacturer’s protocol. Briefly, 50,000 cells per biological replicate were combined with an equal number of Drosophila spike-in cells (Active Motif, 53154) and permeabilized using the kit-supplied lysis buffer containing 0.1% Tween-20 and 0.01% digitonin. Open chromatin regions were subsequently tagmented in vitro using the preassembled Tn5 transposase complex. Genomic DNA was then purified and amplified by PCR to generate sequencing libraries, which were sequenced on an Illumina NovaSeq X platform.

### CUT&Tag

CUT&Tag analysis was performed using the Hyperactive Universal CUT&Tag Assay Kit for Illumina (Vazyme, TD903) according to the manufacturer’s instructions. Briefly, 1 × 10^5^ cells were used for each biological replicate, washed with 1 × wash buffer, and bound to concanavalin A–coated magnetic beads. Cells were permeabilized in digitonin-containing buffer and sequentially incubated with primary antibodies (anti-H3K27me3, anti-H3K36me3, anti-H3K27ac, or normal rabbit IgG) and the corresponding secondary antibody to allow formation of the immunocomplex. DNA tagmentation was then carried out using the preassembled protein A–Tn5 transposome. Drosophila spike-in chromatin (Active Motif, 53083) was added at an equal amount to each reaction for normalization. Following tagmentation, cells were subjected to proteinase K digestion to release DNA fragments, which were subsequently amplified with indexed primers to generate sequencing libraries. Resulting libraries were either analyzed by subjected to high-throughput sequencing on an Illumina NovaSeq X platform.

### caRNA isolation

Briefly, cells were washed in ice-cold PBS supplemented with 1 mM EDTA, pelleted, and lysed on ice in NP-40 lysis buffer (10 mM Tris-HCl, pH 7.5; 150 mM NaCl; 0.05% NP-40). Lysates were gently layered onto a 24% sucrose cushion and centrifuged at 4 °C to separate the cytoplasmic supernatant from the nuclear pellet. The nuclear pellet was rinsed with cold PBS/EDTA, resuspended in glycerol buffer (20 mM Tris-HCl, pH 7.9; 75 mM NaCl; 0.5 mM EDTA; 0.85 mM DTT; 0.125 mM PMSF; 50% glycerol), and extracted with an equal volume of nuclei lysis buffer (10 mM HEPES, pH 7.6; 7.5 mM MgCl₂; 0.2 mM EDTA; 0.3 M NaCl; 1 M urea; 1% NP-40) by brief vortexing. After a short incubation on ice, samples were centrifuged at 4 °C to collect the soluble nucleoplasmic fraction in the supernatant, leaving the chromatin pellet. The chromatin pellet was washed with cold PBS/EDTA and treated with RNase-free DNase I at 37 °C. Chromatin-associated RNA was then extracted from the chromatin fraction using TRIzol and resuspended in RNase-free water.

### In vitro splicing assay

The exon junction complex (EJC)-based in vitro splicing assay was performed as previously described^49^. Briefly, 20 µl splicing reactions were assembled using a Biomek FX (Beckman Coulter) and contained 1× SP buffer (0.5 mM ATP, 20 mM creatine phosphate and 1.6 mM MgCl_2_), 80 µg whole-cell splicing extract, 40 nM biotin-labelled Ad2ΔIVS pre-mRNA and RNasin (1 U µl^−1^). ASOs were added at the indicated concentrations, and reactions were incubated for 90 min at 30 °C. Reactions were diluted in HNT buffer (20 mM HEPES–KOH, pH 7.9, 150 mM NaCl and 0.5% Triton X-100) and captured on black NeutrAvidin-coated plates (Pierce). Bound complexes were detected using anti-eIF4A3 (Santa Cruz Biotechnology, 3F1; 1:350) followed by HRP-conjugated anti-mouse IgG (1:10,000) and chemiluminescent readout (SuperSignal ELISA Femto, Pierce) on an EnVision plate reader (PerkinElmer).

### caRNA-seq

caRNA was isolated from FACS-sorted leukemic cells collected from mice. Ribosomal RNA was depleted from the caRNA using the RiboMinus Eukaryote System v2 (Invitrogen, A15026). Strand-specific libraries were prepared with the SMARTer Stranded Total RNA-Seq Kit v2—Pico Input Mammalian (TaKaRa Bio, 634411) following the manufacturer’s instructions. Three independent biological replicates were processed per condition, and libraries were sequenced on an Illumina NovaSeq X platform.

### mRNA-seq

Total RNA was extracted using standard procedures and treated with DNase I to remove residual genomic DNA. Poly(A)+ mRNA was purifiedi purified from total RNA using the Dynabeads mRNA DIRECT Kit (Invitrogen, 61012) according to the manufacturer’s instructions. Strand-specific libraries were prepared from purified mRNA using the SMARTer Stranded Total RNA-Seq Kit v2—Pico Input Mammalian (TaKaRa Bio, 634411) following the manufacturer’s protocol. Libraries were sequenced on an Illumina NovaSeq X platform.

### ASO and Plasmid Transfection

Antisense oligonucleotides (ASOs; Integrated DNA Technologies) were fully modified with 2′-O-methoxyethyl (2′-MOE) nucleotides and phosphorothioate linkages, and were labeled at the 3′ end with the fluorescent dye FAM to monitor transfection efficiency. The NC5 ASO, targeting non-conserved sequences in the human or mouse genome, was used as a negative control. ASOs and plasmids were transfected into cells using Lipofectamine RNAiMAX (Invitrogen, 13778075) according to the manufacturer’s instructions. All ASO sequences used in this study are provided in **Supplementary Table 1**.

### CLIP-seq

Cultured K562 (WT and SF3B1⁺/⁻) cells were irradiated twice at 254 nm using a Stratalinker (Stratagene) at a total dose of 3,000 J m^−2^ and immediately snap-frozen in liquid nitrogen. Cell pellets were thawed on ice and resuspended in three volumes of ice-cold CLIP lysis buffer (50 mM HEPES pH 7.5, 150 mM KCl, 2 mM EDTA, 0.5% NP-40, 0.5 mM DTT, 1× Halt protease/phosphatase inhibitor cocktail, and 1× RNaseOUT). The cell suspension was passed through a 26-gauge needle to shear genomic DNA, followed by rotation at 4 °C for 15 min. Lysates were then sonicated with a Bioruptor (Diagenode) for five cycles of 30 s on/30 s off. After centrifugation at 21,000 g for 15 min at 4 °C, the cleared supernatant was collected for immunoprecipitation. Supernatants were incubated overnight at 4 °C with Protein A magnetic beads (Abcam, ab205606; Invitrogen, 1001D) pre-bound to anti-Flag antibody. Beads were washed five times with 1 mL of wash buffer (50 mM HEPES pH 7.5, 300 mM KCl, 0.05% NP-40, 1× protease/phosphatase inhibitor cocktail, and 1× RNaseOUT) at 4 °C. Protein–RNA complexes bound to the beads were then partially digested with RNase T1 (8 U μL⁻¹; Thermo Fisher, EN0541) at 22 °C for 10 min with gentle agitation; input samples were processed in parallel under the same conditions. Immunoprecipitated and input samples were resolved by SDS–PAGE, and the corresponding gel fragments were excised and digested with proteinase K (Invitrogen, 25530049) to release RNA. RNA was extracted using TRIzol reagent (Invitrogen, 15596026), end-repaired with T4 PNK (Thermo Fisher, EK0031), and used for library construction with the NEBNext Small RNA Library Prep Set for Illumina (NEB, E7330S). Final libraries were pooled and sequenced on an Illumina NovaSeq X platform.

### Electrophoretic Mobility Shift Assay (EMSA)

Increasing concentrations of purified protein were incubated with 100 nM oligonucleotide probe in 1× binding buffer (20 mM HEPES pH 7.5, 40 mM KCl, 10 mM MgCl₂, 0.1% Triton X-100, 10% glycerol, and 1× RNaseOUT ribonuclease inhibitor [Invitrogen, 10777019]) for 30 min on ice to allow formation of RNA–protein complexes. Reaction mixtures were then loaded onto 10% Novex TBE polyacrylamide gels (Invitrogen, EC62755BOX) and electrophoresed in 0.5× TBE buffer at 4 °C for approximately 2 h. Following electrophoresis, gels were washed twice in 0.5× TBE buffer for 5 min each and stained with SYBR™ Gold Nucleic Acid Gel Stain. Binding curves were fitted using the equation Y = [P] / (K_D_ + [P]), where Y represents the fraction of probe bound at a given protein concentration, calculated from background-subtracted band intensities as bound / (bound + unbound), and [P] is the protein concentration. The fitted K_D_ value was reported as the apparent dissociation constant.

### Fluorescence Microscopy

For immunofluorescence staining, cells were fixed with 4% PFA in DPBS at 37 °C for 5 min, permeabilized with methanol at −20 °C for 8 min, and air-dried at room temperature for 10 min. Samples were washed three times with DPBS and blocked for 1 h at room temperature in blocking buffer (DPBS containing 0.5% BSA, 0.05% Triton X-100, and 1:100 SUPERase·In [Invitrogen, AM2694]). Primary antibodies were diluted in the same blocking buffer at the manufacturer-recommended ratio and incubated with cells for 1 h at room temperature. After three washes with DPBS containing 0.05% Triton X-100, samples were incubated with goat anti-rabbit IgG–AF568 secondary antibody (1:1000; Invitrogen, A-11011) for 1 h at room temperature. Following three additional washes, cells were post-fixed with 4% PFA in DPBS for 30 min at room temperature and rinsed three times with DPBS. Nuclei were counterstained with 2 μg mL⁻¹ Hoechst 33342 (Abcam, ab145597) for 20 min at room temperature, followed by three DPBS washes. Prepared chambers were stored at 4 °C until imaging, which was performed using a Leica SP8 laser-scanning confocal microscope.

### RT–qPCR

To quantify transcript levels, total RNA was reverse transcribed using PrimeScript RT Master Mix (TaKaRa, RR0361) with a combination of oligo(dT) and random hexamer primers. The resulting cDNA was subjected to real-time quantitative PCR using the FastStart Essential DNA Green Master (Roche, 06402712001) and gene-specific primers on a LightCycler 96 system. Relative gene expression levels were calculated using the ΔΔCt method. All primers used in this study are provided in **Supplementary Table 2**.

### Western Blotting

Cells were lysed in RIPA buffer (Thermo Fisher Scientific, 89900) supplemented with 1× Halt protease and phosphatase inhibitor cocktail (Thermo Fisher Scientific, 78441). Protein concentrations were measured using a NanoDrop 8000 spectrophotometer (Thermo Fisher Scientific). Equal amounts of protein were denatured in 1× Laemmli sample buffer (Bio-Rad, 1610747) at 90 °C for 10 min, loaded onto 4–12% NuPAGE Bis-Tris gels (Invitrogen, NP0335BOX), and transferred to PVDF membranes (Thermo Fisher Scientific, 88585) after electrophoresis. Membranes were blocked for 30 min at room temperature in TBST (0.1% Tween-20) containing 5% BSA (Millipore-Sigma, A7030), followed by overnight incubation at 4 °C with appropriately diluted primary antibodies. After washing in TBST, membranes were incubated with HRP-conjugated secondary antibodies for 1 h at room temperature. Protein signals were detected using the SuperSignal West Dura Extended Duration Substrate (Thermo Fisher Scientific, 34075) and imaged on a FluoroChem R system (ProteinSimple). Band intensities were quantified using the Analyze Gel module in Fiji (ImageJ). All antibodies used in this study are provided in **Supplementary Table 3**.

### Hematoxylin and eosin (H&E) staining

For H&E analysis, tissues were harvested and fixed in 10% neutral-buffered formalin, embedded in paraffin, sectioned at 4 µm thickness, and subjected to H&E staining according to standard protocols. Stained sections were examined independently by two board-certified pathologists in a blinded manner to assess the presence of tissue injury, inflammatory infiltration, or other pathological alterations.

### R-loop Dot Blot Assay

Cells were collected following PBS washes, and nuclei were isolated through low-speed centrifugation and lysis. Nuclear lysates were treated with proteinase K, and genomic DNA (retaining RNA–DNA hybrids) was purified by phenol–chloroform extraction and ethanol precipitation. Purified nucleic acids were spotted onto positively charged nylon membranes at varying concentrations and UV-crosslinked. Membranes were probed with either the S9.6 antibody or an anti-dsDNA antibody. After blocking in 5% non-fat milk in TBST, membranes were incubated overnight at 4 °C with primary antibodies, followed by incubation with HRP-conjugated secondary antibodies and chemiluminescent detection. To validate signal specificity, parallel samples were treated with RNase H, RNase T1, or RNase III. Pre-annealed RNA–DNA hybrids, dsDNA, and dsRNA oligonucleotides served as additional controls. Images were quantified using ImageJ, and S9.6 signal intensities were normalized to the corresponding dsDNA signals. The S9.6/dsDNA ratio and standard error were calculated from at least three independent experiments.

### Recombinant Protein Purification

Full-length SETD2 protein was purified according to previously published protocols^51^. For truncated SETD2 constructs, standard molecular cloning strategies were used to generate expression plasmids encoding C-terminal SF3B1–6×His–tagged domains. The human SF3B1 coding sequence was obtained from Origene and amplified using PrimeSTAR GXL DNA polymerase (TaKaRa, R050B) for subcloning. E. coli BL21(DE3) cells were grown in LB medium to an OD_600_ of approximately 0.6, induced with 0.6 mM IPTG at 16 °C for 20 h, and harvested by centrifugation. For purification of MBP-tagged SF3B1 domains, bacterial pellets were resuspended in lysis buffer (25 mM Tris-HCl pH 7.5, 500 mM NaCl, 20 mM imidazole, 10 mM β-mercaptoethanol, and EDTA-free protease inhibitor cocktail [Millipore-Sigma, 4693132001]) and lysed by sonication for 3 min. Lysates were clarified by centrifugation at 26,000 g for 30 min, and the supernatant was applied to Ni²⁺–NTA resin (Thermo Fisher, 88221). After extensive washing with lysis buffer, bound proteins were eluted using the same buffer containing 250 mM imidazole. The eluates were subsequently incubated with amylose resin (NEB, E8021S), and MBP-tagged fusion proteins were eluted with lysis buffer supplemented with 1% maltose. Protein purity was assessed by SDS–PAGE, and eluates were concentrated using Amicon Ultra-15 centrifugal filters. Final protein preparations were aliquoted, snap-frozen in liquid nitrogen, and stored at −80 °C.

### snRNA-KARR-seq Data Analysis

snRNA-KARR-seq data were processed following established protocols with specific modifications tailored for snRNA-target interaction analysis. Specifically, adapter sequences were removed from raw reads, and overlapping paired-end reads (2 × 150 bp) were merged into single-end reads and de-duplicated using fastp. The resulting single-end FASTQ files were first mapped to rDNA sequences (GenBank: KY962518.1) to remove rRNA-derived reads. Unmapped reads were subsequently aligned to the hg38 human genome using STAR. Chimeric reads were extracted from the resulting BAM files and reads containing snRNAs were filtered based on GENCODE annotations. The coordinate pairs (snRNA and the target) were then subjected to duplex group (DG) clustering using a spectral clustering algorithm. Clusters supported by at least one reproducible chimeric read from both replicates were retained. Overlapping regions across replicates were used to define the final interaction coordinates. SnoRNA targets were annotated using the annotatePeaks.pl tool in the HOMER suite. Quantification of snoRNA reads in both unenriched and snRNA-enriched datasets was performed with featureCounts, followed by differential abundance analysis using DESeq2. The secondary structure and predicted free energy of snoRNA–target duplexes were modeled using RNAplex (parameters: -c 40 -l 40) and visualized using VARNA. Binding regions of snoRNAs were visualized by converting BED files into BigWig format via bedGraphToBigWig, and genome-wide coverage tracks were generated using CoolBox. Motif enrichment analysis of U1 targets was carried out using findMotifs.pl in the HOMER suite, with background sequences generated by the shuffle function in BEDTools.

### ATAC-seq Data Analysis

ATAC-seq data were processed following the workflow described by Reske et al., with the complete code available at https://github.com/reskejak/ATAC-seq. Briefly, sequencing reads were trimmed, aligned to the hg38 and BDGP6 reference genome, and filtered for duplicate and mitochondrial reads as described. Peaks were called for each sample independently, and a peak set was generated by merging all peaks across the experimental batch using BEDTools. Read counts were quantified over the merged peak set to obtain a uniform measure of chromatin accessibility across samples. To enable accurate comparison of global chromatin accessibility, read counts were normalized using the Drosophila melanogaster chromatin spike-in included during library preparation.

### WGS Data Analysis

WGS reads were trimmed using fastp and aligned to the hg38 reference genome with BWA-MEM using sample-specific read group information. Aligned reads were converted to BAM format and sorted with samtools. Genome-wide coverage was quantified by generating bigWig files for each sample and computing bin-level read counts using multiBigwigSummary (deepTools) with a 100-kb bin size. The resulting per-bin count matrix was used to estimate relative copy-number changes. For each genomic bin, read counts from experimental samples were normalized to the mean coverage of the control group, and copy-number ratios were calculated using a custom script to generate genome-wide CNV profiles.

### CUT&Tag and CUT&Run Data Analysis

CUT&Tag and CUT&Run sequencing reads were trimmed using fastp and aligned to the human reference genome (hg38) with Bowtie2 in paired-end mode. Resulting alignments were converted to BAM format, sorted, and filtered to remove unmapped and mitochondrial reads. In parallel, the same alignment workflow was performed against the Drosophila melanogaster BDGP6 genome to quantify the spike-in chromatin included for normalization. For each sample, human and spike-in BAM files were indexed and mapping statistics were generated using samtools. Peaks were called using MACS3 in broad-peak mode with the following parameters: -f BAMPE, --broad, --broad-cutoff 0.1, -q 0.05, --keep-dup all, and --nolambda, and a peak set was generated by merging all peaks across the experimental batch using BEDTools. To allow accurate comparison of global histone-modification levels across samples, read counts were normalized to the number of mapped, non-mitochondrial Drosophila spike-in reads.

### RNA-seq Data Analysis

RNA-seq sequencing reads were first subjected to adapter trimming and quality filtering using Cutadapt (vX.X) with the following parameters: -j 24 -e 0.1 -n 1 -O 1 -q 20 -m 29 --nextseq-trim=20. To handle specific library architectures, 3 bp were removed from the 5’ end of the R2 reads (-U 3), and the adapter sequences AGATCGGAAGAGC and NNNAGATCGGAAGAGC were utilized for R1 and R2, respectively. To ensure the accuracy of transcriptomic quantification, trimmed reads were first aligned to the human ribosomal RNA (rRNA) reference using HISAT2. Reads that did not map to rRNA were subsequently aligned to the human reference genome (hg38) using HISAT2. The resulting alignments were converted to BAM format, sorted, and indexed using samtools. For gene-level quantification, the number of reads mapped to each gene was calculated using featureCounts (from the Subread package) against the GENCODE v45 (hg38) annotation GTF file. Differential gene expression analysis was performed using DESeq2 with default parameters. Significant differentially expressed genes (DEGs) were identified based on the built-in statistical framework of DESeq2, providing a basis for downstream functional enrichment and pathway analysis.

### CLIP-seq Data Analysis

CLIP-seq reads were trimmed using cutadapt (parameters: -e 0.1, -q 20, -m 29, --nextseq-trim=20, -U 3, -a AGATCGGAAGAGC, -A GATCGTCGGACTGTAGA) to remove adapters and low-quality bases. Trimmed reads1 (R1) were aligned to the hg38 reference genome using STAR with the following settings: --alignEndsType EndToEnd, --quantMode TranscriptomeSAM, --outFilterMultimapNmax 20, --outFilterMatchNmin 24, and --alignMatesGapMax 15000.

Alignments were output as coordinate-sorted BAM files. To obtain strand-specific binding signals, mapped reads were separated by strand using samtools: forward-strand reads with -F 16 and reverse-strand reads with -f 16. Peak calling was performed separately for each strand using MACS3 (-f BAM, --nomodel, --extsize 30, --keep-dup all, -g hs) by comparing immunoprecipitated (IP) samples against their matched input controls. Resulting peak sets provided strand-resolved binding profiles. To quantify RNA binding strength, read counts were obtained for both IP and input samples using featureCounts at the gene level. Counts were normalized using CPM, and enrichment ratios were computed for each gene in each sample. Differential binding between WT and K700E groups was assessed using limma for statistical test, focusing on RNA genes shorter than 200 nt to specifically evaluate small-RNA binding events.

### Quantification and Statistical Analysis

Unless otherwise stated, data are presented as mean ± standard deviation (SD). Comparisons between two groups were performed using two-tailed unpaired Student’s t-tests, whereas comparisons among multiple groups were conducted using one-way ANOVA. One-way ANOVA was used for analyses of tumor weight and toxicology-related measurements. Two-way ANOVA was applied to time-course experiments, including in vitro cell viability assays and tumor growth curves. Survival analyses were performed using the log-rank (Mantel–Cox) test. Statistical significance was defined as p < 0.05. Data processing, statistical analyses, and visualization were carried out using GraphPad Prism v10 (GraphPad Software) and SPSS v20 (IBM SPSS Statistics). For box-and-whisker plots, the center line represents the median, box boundaries correspond to the upper and lower quartiles, and whiskers extend to 1.5× the interquartile range.

## Acknowledgements

We are grateful to Dr. Pieter W. Faber for assistance with high-throughput sequencing and to the University of Chicago Human Tissue Resource Center (RRID: SCR_019199) and the Integrated Small Animal Imaging Research Resource (RRID: SCR_017923) for technical support with mouse imaging and data analysis. This work is supported by the National Institute of Health RM1 HG008935 and R01 ES030546 to C.H.; C.H. is an investigator of the Howard Hughes Medical Institute. O.A.W. is supported by the Neil S. Hirsch Foundation, Damon Runyon Foundation, Edward P. Evans Foundation, BreakThrough Cancer, and the National Institutes of Health (R35CA304457, R01 CA251138, CA283364, CA242020, HL128239, P50 CA254838-01, P30 CA08748). R.K.B. was supported in part by the NIH/NCI (R01 CA251138), NIH/NHLBI (R01 HL128239, R01 HL151651) and the Blood Cancer Discoveries Grant program through the Leukemia & Lymphoma Society, Mark Foundation for Cancer Research, and Paul G. Allen Frontiers Group (8023-20). R.K.B holds the McIlwain Family Endowed Chair in Data Science.

## Author contributions statement

C.H. supervised the entire project. C.H., P.X., H.L., B.L., R.K.B., and O.A.-W. conceived the original idea and designed the original studies. P.X. performed most cell line experiments with the help from YY.J., CW.J., YQ.P., J.M., LJ.Z., XH.Z., YX.A., and JB.W.. H.L. performed bioinformatics analysis and interpreted the data with the help of B.L.. P.X., and YQ.P. performed *in vivo* studies with mice and primary cells. P.X., H.L., B.L., LL.W., R.K.B., O.A.-W., and C.H. wrote the manuscript. All authors approved the final manuscript.

## Competing interests

C.H. is a scientific founder, a member of the scientific advisory board, and equity holder of Aferna Bio, Inc., AllyRNA, Inc. and Ellis Bio Inc., a scientific cofounder and equity holder of Accent Therapeutics, Inc., and a member of the scientific advisory board of Rona Therapeutics and Element Biosciences. O.A.-W. has served as a consultant for Amphista Therapeutics, MagnetBio, Volastra, Minovia, and Astra Zeneca, and is on the Scientific Advisory Board of Envisagenics Inc., Harmonic Discovery Inc., and Pfizer Boulder; O.A.-W. received research funding from Nurix Therapeutics as well as research funding from Minovia Therapeutics and Astra Zeneca unrelated to the current manuscript. The other authors declare no competing interests. R.K.B. is a founder and scientific advisor of Codify Therapeutics and Synthesize Bio and holds equity in both companies. R.K.B. has received research funding from Codify Therapeutics unrelated to the current work.

## Data availability

All data are available in the Article and Supplementary Information. All high-throughput sequencing data generated in this study have been deposited in the NCBI Gene Expression Omnibus (GEO) under accession numbers GSE312499, GSE312770, GSE312771, GSE312851, GSE312852, GSE312853, GSE312856, GSE317021, GSE312858, and GSE316375. Source data are provided with this paper. All other data supporting the findings of this study are available from the corresponding author upon reasonable request. The data will be made publicly available upon publication.

## References

1 Bradley, R. K. & Anczukow, O. RNA splicing dysregulation and the hallmarks of cancer. Nat Rev Cancer 23, 135–155 (2023).

2 Kim, E. et al. SRSF2 Mutations Contribute to Myelodysplasia by Mutant-Specific Effects on Exon Recognition. Cancer Cell 27, 617–630 (2015).

3 Quesada, V. et al. Exome sequencing identifies recurrent mutations of the splicing factor SF3B1 gene in chronic lymphocytic leukemia. Nat Genet 44, 47–52 (2011).

4 Bonnal, S., Vigevani, L. & Valcarcel, J. The spliceosome as a target of novel antitumour drugs. Nat Rev Drug Discov 11, 847–859 (2012).

5 Wan, Y. & Wu, C. J. SF3B1 mutations in chronic lymphocytic leukemia. Blood 121, 4627–4634 (2013).

6 Rogalska, M. E., Vivori, C. & Valcarcel, J. Regulation of pre-mRNA splicing: roles in physiology and disease, and therapeutic prospects. Nat Rev Genet 24, 251–269 (2023).

7 Berg, M. G. et al. U1 snRNP determines mRNA length and regulates isoform expression. Cell 150, 53–64 (2012).

8 Wan, R., Bai, R., Zhan, X. & Shi, Y. How Is Precursor Messenger RNA Spliced by the Spliceosome? Annu Rev Biochem 89, 333–358 (2020).

9 Kiss, T. Small nucleolar RNAs: an abundant group of noncoding RNAs with diverse cellular functions. Cell 109, 145–148 (2002).

10 Reichow, S. L., Hamma, T., Ferre-D’Amare, A. R. & Varani, G. The structure and function of small nucleolar ribonucleoproteins. Nucleic Acids Res 35, 1452–1464 (2007).

11 Yin, Y. et al. U1 snRNP regulates chromatin retention of noncoding RNAs. Nature 580, 147–150 (2020).

12 Jobert, L. et al. Human U1 snRNA forms a new chromatin-associated snRNP with TAF15. EMBO Rep 10, 494–500 (2009).

13 Yun, H. et al. The landscape of RNA-chromatin interaction reveals small non-coding RNAs as essential mediators of leukemia maintenance. Leukemia 38, 1688–1698 (2024).

14 Kaida, D. et al. U1 snRNP protects pre-mRNAs from premature cleavage and polyadenylation. Nature 468, 664–668 (2010).

15 So, B. R. et al. A Complex of U1 snRNP with Cleavage and Polyadenylation Factors Controls Telescripting, Regulating mRNA Transcription in Human Cells. Mol Cell 76, 590–599 e594 (2019).

16 Ebert, B. & Bernard, O. A. Mutations in RNA splicing machinery in human cancers. N Engl J Med 365, 2534–2535 (2011).

17 Quesada, V., Ramsay, A. J. & Lopez-Otin, C. Chronic lymphocytic leukemia with SF3B1 mutation. N Engl J Med 366, 2530 (2012).

18 Damm, F., Nguyen-Khac, F., Fontenay, M. & Bernard, O. A. Spliceosome and other novel mutations in chronic lymphocytic leukemia and myeloid malignancies. Leukemia 26, 2027–2031 (2012).

19 Wang, L. et al. SF3B1 and other novel cancer genes in chronic lymphocytic leukemia. N Engl J Med 365, 2497–2506 (2011).

20 Damm, F. et al. Acquired initiating mutations in early hematopoietic cells of CLL patients. Cancer Discov 4, 1088–1101 (2014).

21 Damianov, A., Lin, C. H., Zhang, J., Manley, J. L. & Black, D. L. Cancer-associated SF3B1 mutation K700E causes widespread changes in U2/branchpoint recognition without altering splicing. Proc Natl Acad Sci U S A 122, e2423776122 (2025).

22 Ji, X. et al. SR proteins collaborate with 7SK and promoter-associated nascent RNA to release paused polymerase. Cell 153, 855–868 (2013).

23 Cusan, M. et al. SF3B1 mutation and ATM deletion codrive leukemogenesis via centromeric R-loop dysregulation. J Clin Invest 133 (2023).

24 Bland, P. et al. SF3B1 hotspot mutations confer sensitivity to PARP inhibition by eliciting a defective replication stress response. Nat Genet 55, 1311–1323 (2023).

25 Wu, T. et al. KARR-seq reveals cellular higher-order RNA structures and RNA-RNA interactions. Nat Biotechnol 42, 1909–1920 (2024).

26 Liu, B. et al. snoRNA-facilitated protein secretion revealed by transcriptome-wide snoRNA target identification. Cell 188, 465–483 e422 (2025).

27 Kiss-Laszlo, Z., Henry, Y., Bachellerie, J. P., Caizergues-Ferrer, M. & Kiss, T. Site-specific ribose methylation of preribosomal RNA: a novel function for small nucleolar RNAs. Cell 85, 1077–1088 (1996).

28 Jurica, M. S. & Moore, M. J. Pre-mRNA splicing: awash in a sea of proteins. Mol Cell 12, 5–14 (2003).

29 Pomeranz Krummel, D. A., Oubridge, C., Leung, A. K., Li, J. & Nagai, K. Crystal structure of human spliceosomal U1 snRNP at 5.5 A resolution. Nature 458, 475–480 (2009).

30 Huang, H. et al. Histone H3 trimethylation at lysine 36 guides m(6)A RNA modification co-transcriptionally. Nature 567, 414–419 (2019).

31 Wagner, E. J. & Carpenter, P. B. Understanding the language of Lys36 methylation at histone H3. Nat Rev Mol Cell Biol 13, 115–126 (2012).

32 Carvalho, S. et al. Histone methyltransferase SETD2 coordinates FACT recruitment with nucleosome dynamics during transcription. Nucleic Acids Res 41, 2881–2893 (2013).

33 Carrozza, M. J. et al. Histone H3 methylation by Set2 directs deacetylation of coding regions by Rpd3S to suppress spurious intragenic transcription. Cell 123, 581–592 (2005).

34 Keogh, M. C. et al. Cotranscriptional set2 methylation of histone H3 lysine 36 recruits a repressive Rpd3 complex. Cell 123, 593–605 (2005).

35 Cookis, T., Lydecker, A., Sauer, P., Kasinath, V. & Nogales, E. Structural basis for the inhibition of PRC2 by active transcription histone posttranslational modifications. Nat Struct Mol Biol 32, 393–404 (2025).

36 Yuan, H. et al. SETD2 Restricts Prostate Cancer Metastasis by Integrating EZH2 and AMPK Signaling Pathways. Cancer Cell 38, 350–365 e357 (2020).

37 Schmitges, F. W. et al. Histone methylation by PRC2 is inhibited by active chromatin marks. Mol Cell 42, 330–341 (2011).

38 Markert, J. W., Soffers, J. H. & Farnung, L. Structural basis of H3K36 trimethylation by SETD2 during chromatin transcription. Science 387, 528–533 (2025).

39 Sharma, S., Wongpalee, S. P., Vashisht, A., Wohlschlegel, J. A. & Black, D. L. Stem-loop 4 of U1 snRNA is essential for splicing and interacts with the U2 snRNP-specific SF3A1 protein during spliceosome assembly. Genes Dev 28, 2518–2531 (2014).

40 Inoue, D. et al. Spliceosomal disruption of the non-canonical BAF complex in cancer. Nature 574, 432–436 (2019).

41 Boddu, P. C. et al. Transcription elongation defects link oncogenic SF3B1 mutations to targetable alterations in chromatin landscape. Mol Cell 84, 1475–1495 e1418 (2024).

42 Singh, S. et al. SF3B1 mutations induce R-loop accumulation and DNA damage in MDS and leukemia cells with therapeutic implications. Leukemia 34, 2525–2530 (2020).

43 Landau, D. A. et al. Evolution and impact of subclonal mutations in chronic lymphocytic leukemia. Cell 152, 714–726 (2013).

44 Pawlik, P. et al. Clone copy number diversity is linked to survival in lung cancer. Nature 646, 190–197 (2025).

45 Nadeu, F. et al. Clinical impact of clonal and subclonal TP53, SF3B1, BIRC3, NOTCH1, and ATM mutations in chronic lymphocytic leukemia. Blood 127, 2122–2130 (2016).

46 Desterro, J., Bak-Gordon, P. & Carmo-Fonseca, M. Targeting mRNA processing as an anticancer strategy. Nat Rev Drug Discov 19, 112–129 (2020).

47 Wilkinson, M. E., Charenton, C. & Nagai, K. RNA Splicing by the Spliceosome. Annu Rev Biochem 89, 359–388 (2020).

48 Bonnal, S. C., Lopez-Oreja, I. & Valcarcel, J. Roles and mechanisms of alternative splicing in cancer - implications for care. Nat Rev Clin Oncol 17, 457–474 (2020).

49 Berg, M. G. et al. A quantitative high-throughput in vitro splicing assay identifies inhibitors of spliceosome catalysis. Mol Cell Biol 32, 1271–1283 (2012).

