## Extended Data Figures for "A U1–U3 snRNA–snoRNA interaction couples SF3B1 mutation to chromatin-state rewiring and genome instability"

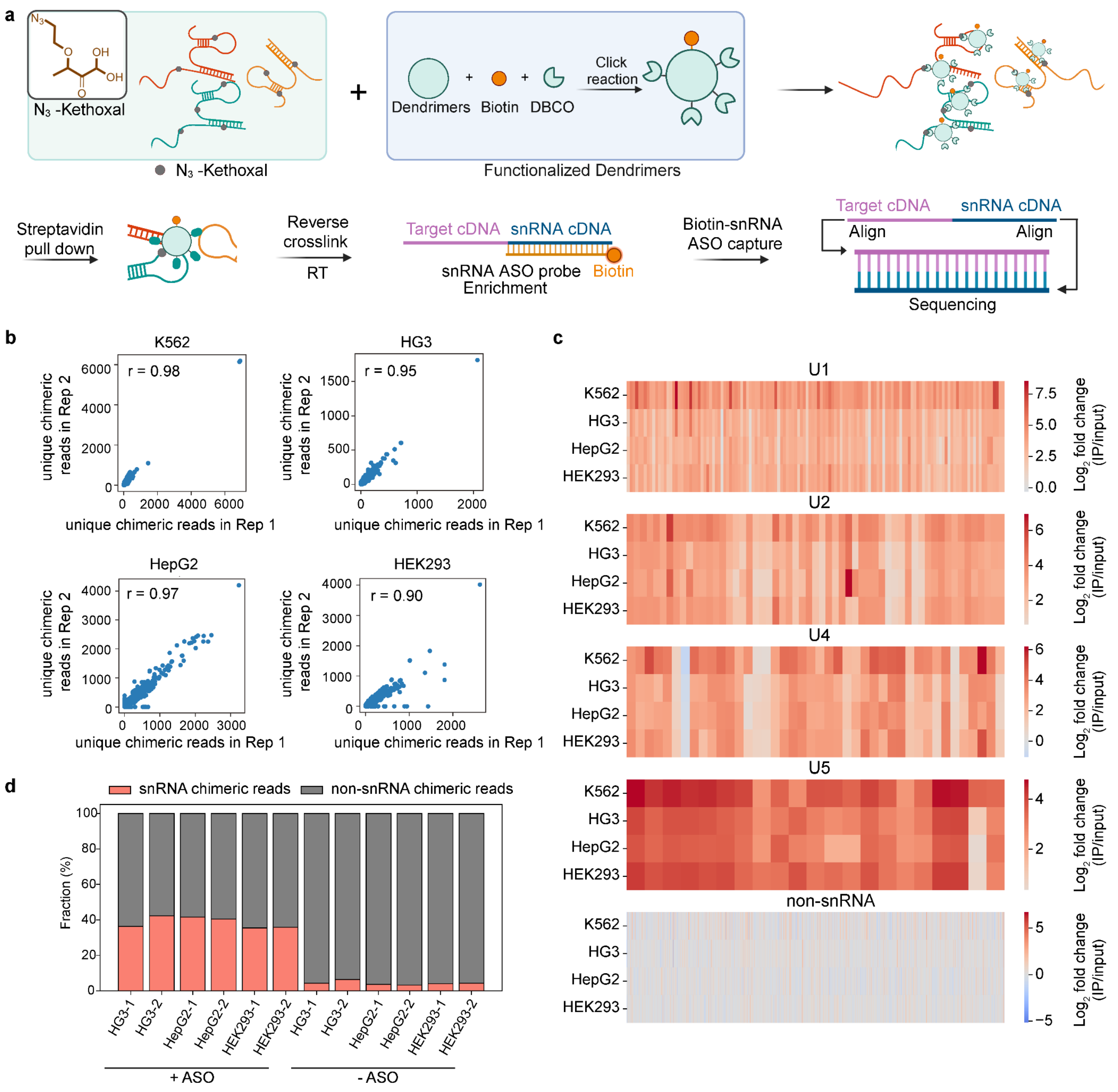


**Extended Data Fig. 1 | snRNA enrichment and binding preferences.**

**a**, Workflow schematic of the snRNA-enrichment KARR-seq (snRNA-KARR-seq) method. **b**, Correlation of unique chimeric reads obtained from biological replicates across K562, HG3, HepG2, and HEK293 cells. **c**, Enrichment efficiency of canonical snRNAs (U1, U2, U4, and U5) and non-snRNA total reads across K562, HG3, HepG2, and HEK293 cells. **d**, Efficiency of enrichment for chimeric reads involving U1, U2, U4, and U5 across K562, HG3, HepG2, and HEK293 cells.

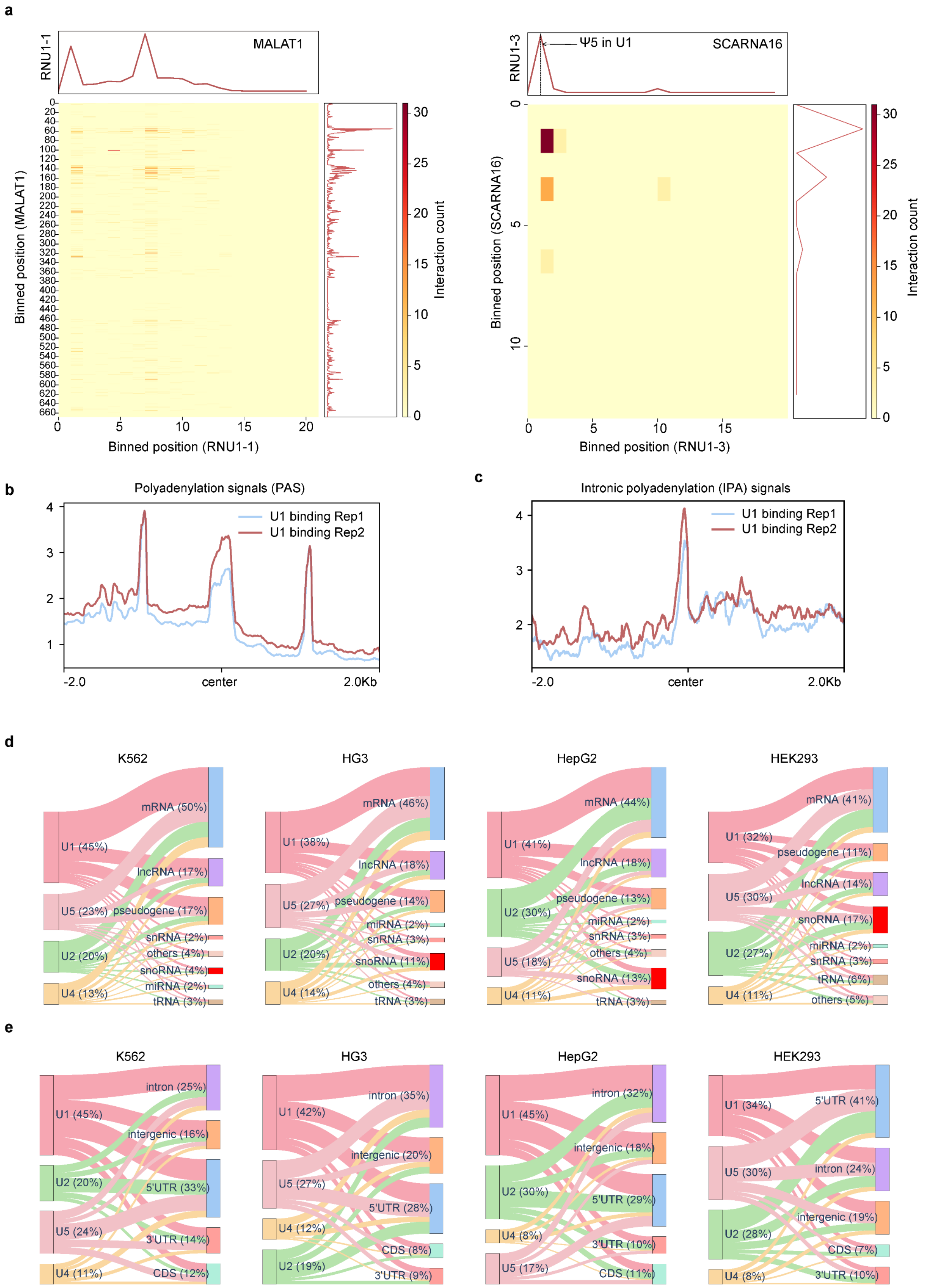
a
RNU1-1
MALAT1
Binned position (MALAT1)
Binned position (RNU1-1)
Interaction count
RNU1-3
Ψ5 in U1
SCARNA16
Binned position (SCARNA16)
Binned position (RNU1-3)
Interaction count
b
Polyadenylation signals (PAS)
U1 binding Rep1
U1 binding Rep2
-2.0
center
2.0Kb
c
Intronic polyadenylation (IPA) signals
U1 binding Rep1
U1 binding Rep2
-2.0
center
2.0Kb
d
K562
U1 (45%)
U5 (23%)
U2 (20%)
U4 (13%)
mRNA (50%)
lncRNA (17%)
pseudogene (17%)
snRNA (2%)
others (4%)
snoRNA (4%)
miRNA (2%)
tRNA (3%)
HG3
U1 (38%)
U5 (27%)
U2 (20%)
U4 (14%)
mRNA (46%)
lncRNA (18%)
pseudogene (14%)
miRNA (2%)
snRNA (3%)
snoRNA (11%)
others (4%)
tRNA (3%)
HepG2
U1 (41%)
U2 (30%)
U5 (18%)
U4 (11%)
mRNA (44%)
lncRNA (18%)
pseudogene (13%)
miRNA (2%)
snRNA (3%)
others (4%)
snoRNA (13%)
tRNA (3%)
HEK293
U1 (32%)
U5 (30%)
U2 (27%)
U4 (11%)
mRNA (41%)
pseudogene (11%)
lncRNA (14%)
snoRNA (17%)
miRNA (2%)
snRNA (3%)
tRNA (6%)
others (5%)
e
K562
U1 (45%)
U2 (20%)
U5 (24%)
U4 (11%)
intron (25%)
intergenic (16%)
5'UTR (33%)
3'UTR (14%)
CDS (12%)
HG3
U1 (42%)
U5 (27%)
U4 (12%)
U2 (19%)
intron (35%)
intergenic (20%)
5'UTR (28%)
CDS (8%)
3'UTR (9%)
HepG2
U1 (45%)
U2 (30%)
U4 (8%)
U5 (17%)
intron (32%)
intergenic (18%)
5'UTR (29%)
3'UTR (10%)
CDS (11%)
HEK293
U1 (34%)
U5 (30%)
U2 (28%)
U4 (8%)
5'UTR (41%)
intron (24%)
intergenic (19%)
CDS (7%)
3'UTR (10%)

**Extended Data Fig. 2 | U1 snRNA enrichment and binding preferences.**

**a**, 2D-interaction heatmaps illustrating known U1–MALAT1 (left) and U1–SCARNA16 (right) RNA interactions in K562 cells. **b**–**c**, U1 snRNA occupancy around polyadenylation signal (PAS) site (**b**) and intronic polyadenylation (IPA) site (**c**) in K562 cells. **d**, Sankey diagrams depicting the distribution of snRNA target classes across K562, HG3, HepG2, and HEK293 cells. **e**, Sankey diagrams showing the proportion and identity of mRNA/pe-mRNA targets across K562, HG3, HepG2, and HEK293 cells.

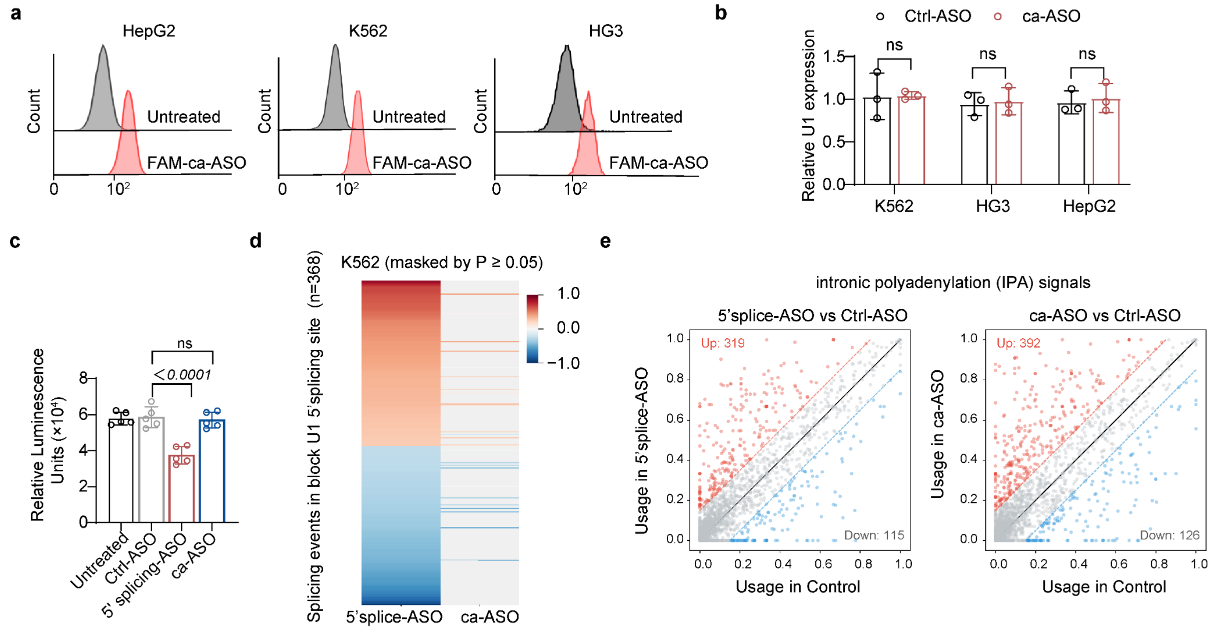


**Extended Data Fig. 3 | Validation of ca-ASO activity and specificity.**

**a**, Flow cytometry confirmed efficient uptake of FAM-labeled ca-ASO in HepG2, K562, and HG3 cells ($n$ = 3 biological per group). **b**, qRT–PCR analysis of U1 snRNA expression in K562, HG3, and HepG2 cells following ca-ASO treatment ($n$ = 3 biological per group). **c**, In vitro splicing of biotin-labelled Ad2ΔIVS pre-mRNA in whole-cell extract in the presence of the indicated ASOs, with exon junction complex (EJC) deposition quantified by HRP-based chemiluminescence ($n$ = 5 biological per group). **d**, Heat map of differential splicing changes in K562 cells following transfection with the indicated ASOs. Events are ordered by direction and magnitude of change; non-significant values are masked ($P \geq 0.05$; $n$ = 3 biological per group). **e**, Scatter plots of changes in intronic polyadenylation (IPA) site usage in K562 cells transfected with the indicated ASOs relative to control conditions ($n$ = 3 biological per group). Data are mean ± s.d for **b** and **c**. *P* value was determined using an unpaired two-tailed *t* test (**b** and **c**). ns indicates no statistically significant difference.

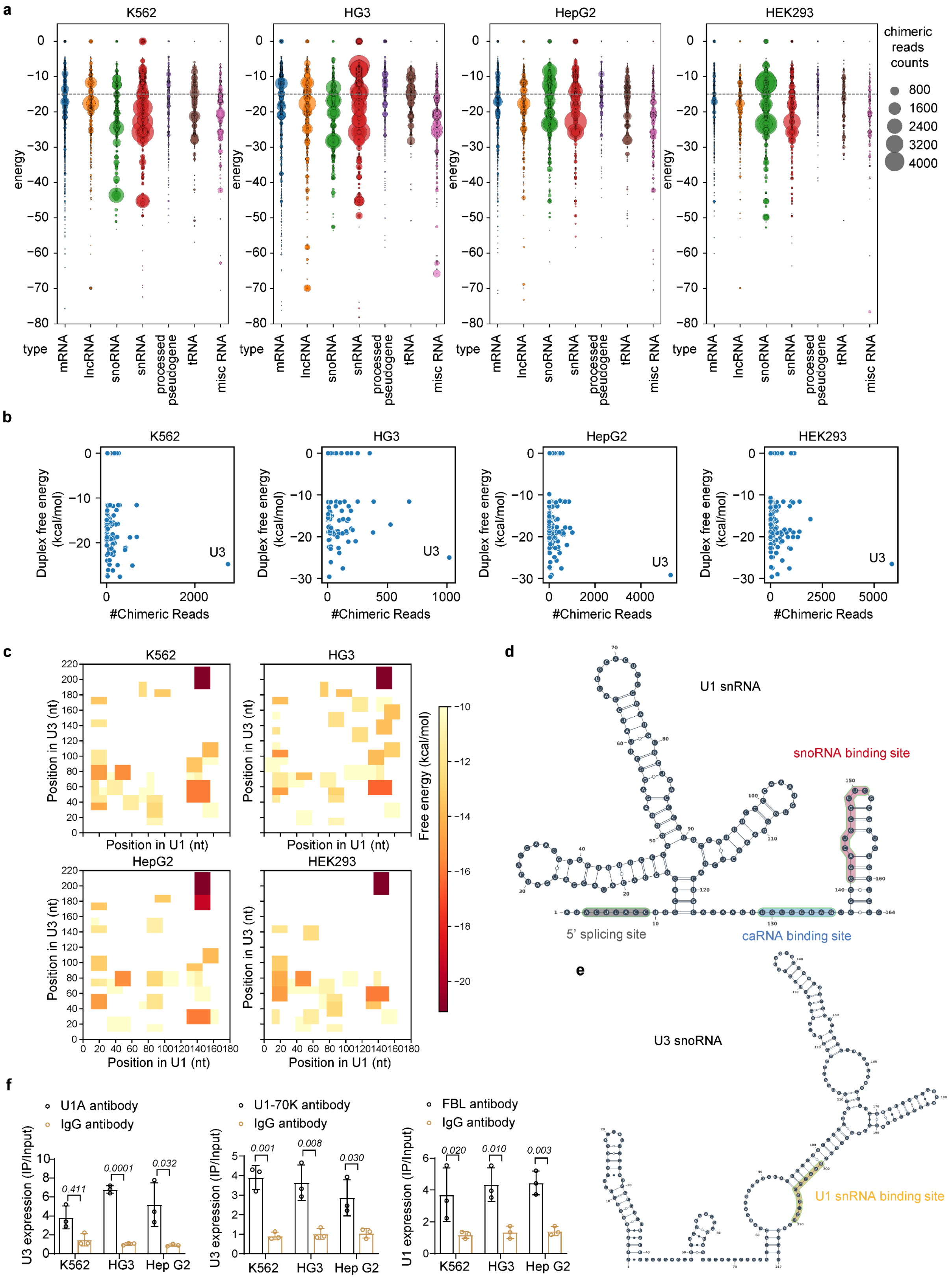
a
K562
HG3
HepG2
HEK293
chimeric reads counts
800
1600
2400
3200
4000
energy
type
mRNA
lncRNA
snoRNA
snRNA
processed pseudogene
tRNA
misc RNA
b
Duplex free energy (kcal/mol)
#Chimeric Reads
U3
c
Position in U3 (nt)
Position in U1 (nt)
Free energy (kcal/mol)
d
U1 snRNA
snoRNA binding site
5' splicing site
caRNA binding site
e
U3 snoRNA
U1 snRNA binding site
f
U1A antibody
IgG antibody
U1-70K antibody
FBL antibody
U3 expression (IP/Input)
U1 expression (IP/Input)
K562
HG3
Hep G2
0.411
0.0001
0.032
0.001
0.008
0.030
0.020
0.010
0.003

**Extended Data Fig. 4 | KARR-seq analysis of snRNA–RNA interactions.**

**a**, Predicted base pairing free energy values of snRNA–target duplexes across K562, HG3, HepG2, and HEK293 cells. **b**, Comparative analysis of chimeric read counts and predicted ΔG values for diverse snRNA-target interactions identified via KARR-seq across K562, HG3, HepG2, and HEK293 cells. **c**, Heatmap displaying predicted ΔG values of U1-U3 interactions across K562, HG3, HepG2, and HEK293 cells. **d**, U1 secondary structure model with color-coded binding motifs: splice site (grey), caRNA binding site (blue), and U1-U3 interaction region (red). **e**, Secondary structure model of U3 snoRNA, highlighting the predicted U1 binding site in yellow. **f**, RIP–qPCR of chromatin fraction from K562, HG3, and HepG2 cells showed enrichment of U3 snoRNA using U1A or U1-70K antibodies compared with IgG controls, and enrichment of U1 snRNA using FBL antibody compared with IgG controls ($n$ = 3 biological per group). Data are mean ± s.d for **f**. $P$ value was determined using an unpaired two-tailed $t$ test (**f**).

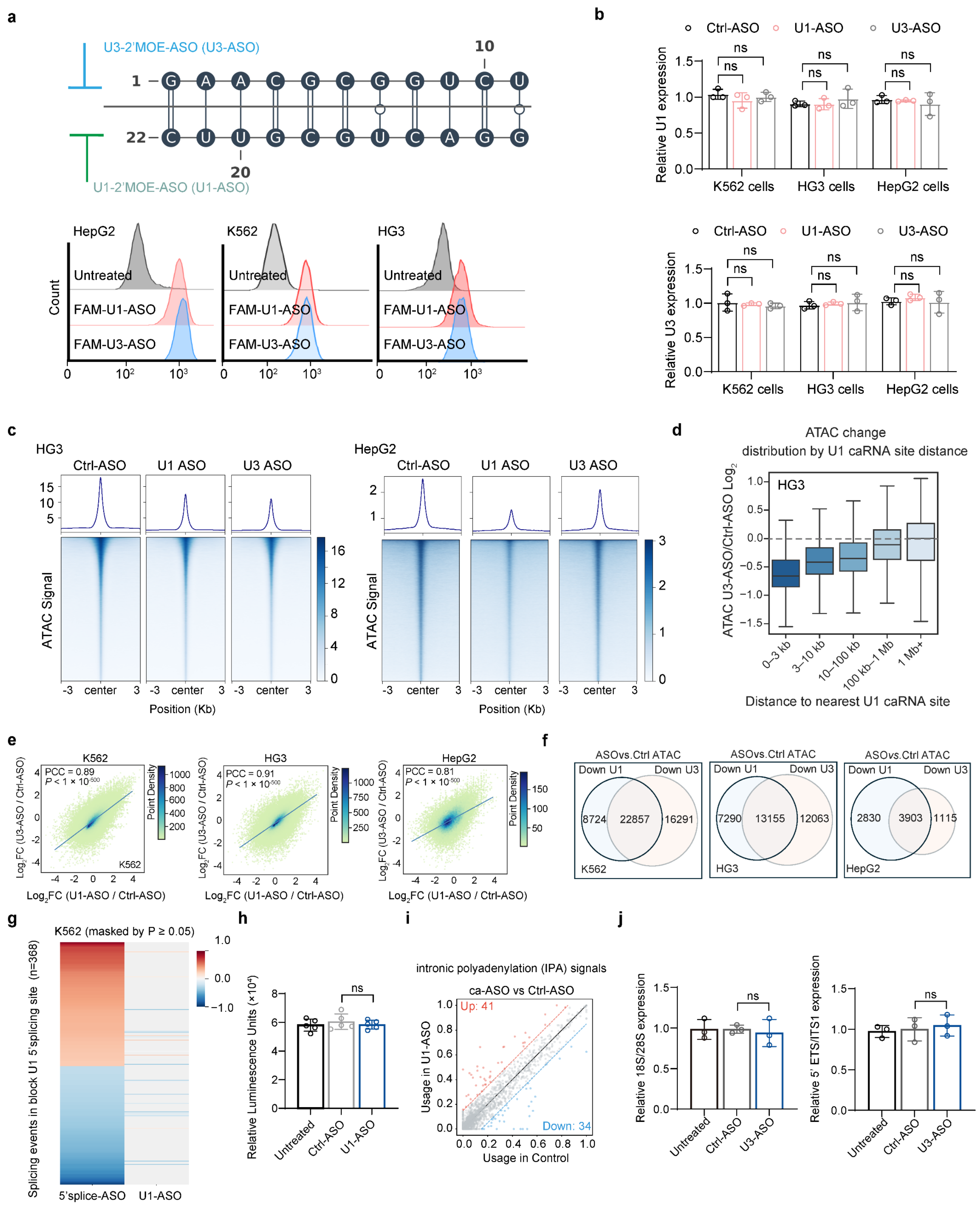

**Extended Data Fig. 5 | Effects of U1-ASO and U3-ASO on chromatin accessibility.**

**a**, Flow cytometry analysis of FAM-labelled ASO uptake in HepG2, K562, and HG3 cells (*n* = 3 biological per group). **b**, RT–qPCR quantification of U1 snRNA and U3 snoRNA abundance following ASO treatment in HepG2, K562, and HG3 cells (*n* = 3 biological per group). **c**, Average ATAC-seq profiles and heat maps of chromatin accessibility in K562 and HG3 cells following U1-ASO or U3-ASO treatment. **d**, ATAC-seq signal changes as a function of distance from U1–caRNA binding sites. **e**, Correlation analysis of differential ATAC-seq peaks following U1-ASO or U3-ASO treatment in K562, HG3, and HepG2 cells. **f**, Venn diagram showing the overlap of differential ATAC peaks in K562, HG3, and HepG2 cells. **g**, Heat map of differential splicing changes in K562 cells following transfection with the indicated ASOs. Events are ordered by direction and magnitude of change; non-significant values are masked ($P \geq 0.05$; *n* = 3 biological per group). **h**, In vitro splicing of biotin-labelled Ad2ΔIVS pre-mRNA in whole-cell extract in the presence of the indicated ASOs, with exon junction complex (EJC) deposition quantified by HRP-based chemiluminescence (*n* = 5 biological per group). **i**, Scatter plots of changes in intronic polyadenylation (IPA) site usage in K562 cells transfected with the indicated ASOs relative to control conditions (*n* = 3 biological per group). **j**, RT–qPCR quantification of mature rRNA (18S and 28S) and pre-rRNA (5′ETS and ITS1) levels in K562 cells treated with the indicated ASOs (*n* = 3 biological per group). For **d**, the box plots show the median (centre line), upper and lower quartiles (box limits) and 1–99% (whiskers). *P* value was determined using an unpaired two-tailed *t* test (**b**), and Pearson's correlation coefficient (PCC) was used to assess correlation (**e**). ns indicates no statistically significant difference. ns indicates no statistically significant difference.

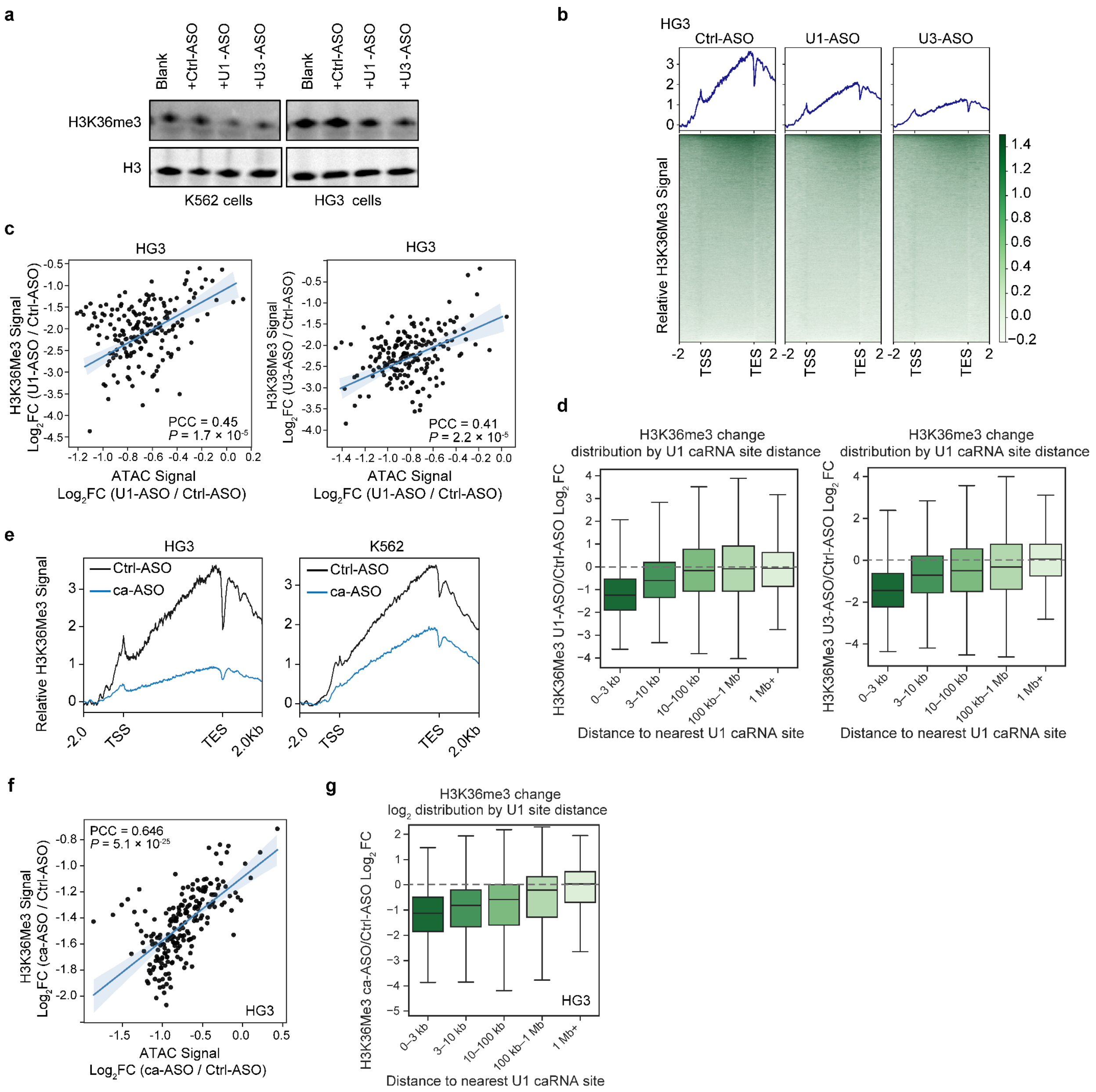


**Extended Data Fig. 6 | Effects of U1-ASO and U3-ASO on histone modification.**

**a**, Western blot analysis of H3K36me3 in K562 and HG3 cells following U1-ASO or U3-ASO treatment. **b**, CUT&Tag profiles showing changes in H3K36me3 peaks in HG3 cells after U1-ASO or U3-ASO treatment. **c**, Scatter plots showing the relationship between H3K36me3 signal changes and ATAC-seq signal changes in HG3 cells after U1–ASO treatment in HG3 cells. **d**, H3K36me3 changes in K562 cells displayed a clear spatial gradient, with the strongest reductions occurring within a few kilobases of U1–caRNA binding sites and progressively diminishing toward zero at more distal regions. **e**, CUT&Tag profiles of H3K36me3 in K562 and HG3 cells following ca-ASO treatment. **f**, Scatter plots showing the relationship between

H3K36me3 signal changes and ATAC-seq signal changes after ca–ASO treatment in HG3 cells. **g**, H3K36me3 changes displayed a clear spatial gradient, with the strongest reductions occurring within a few kilobases of U1–caRNA binding sites and progressively diminishing toward zero at more distal regions in HG3 cells. $n = 3$ biological per group. For **d** and **g**, the box plots show the median (centre line), upper and lower quartiles (box limits) and 1–99% (whiskers). Pearson's correlation coefficient (PCC) was used to assess correlation (**c** and **f**).

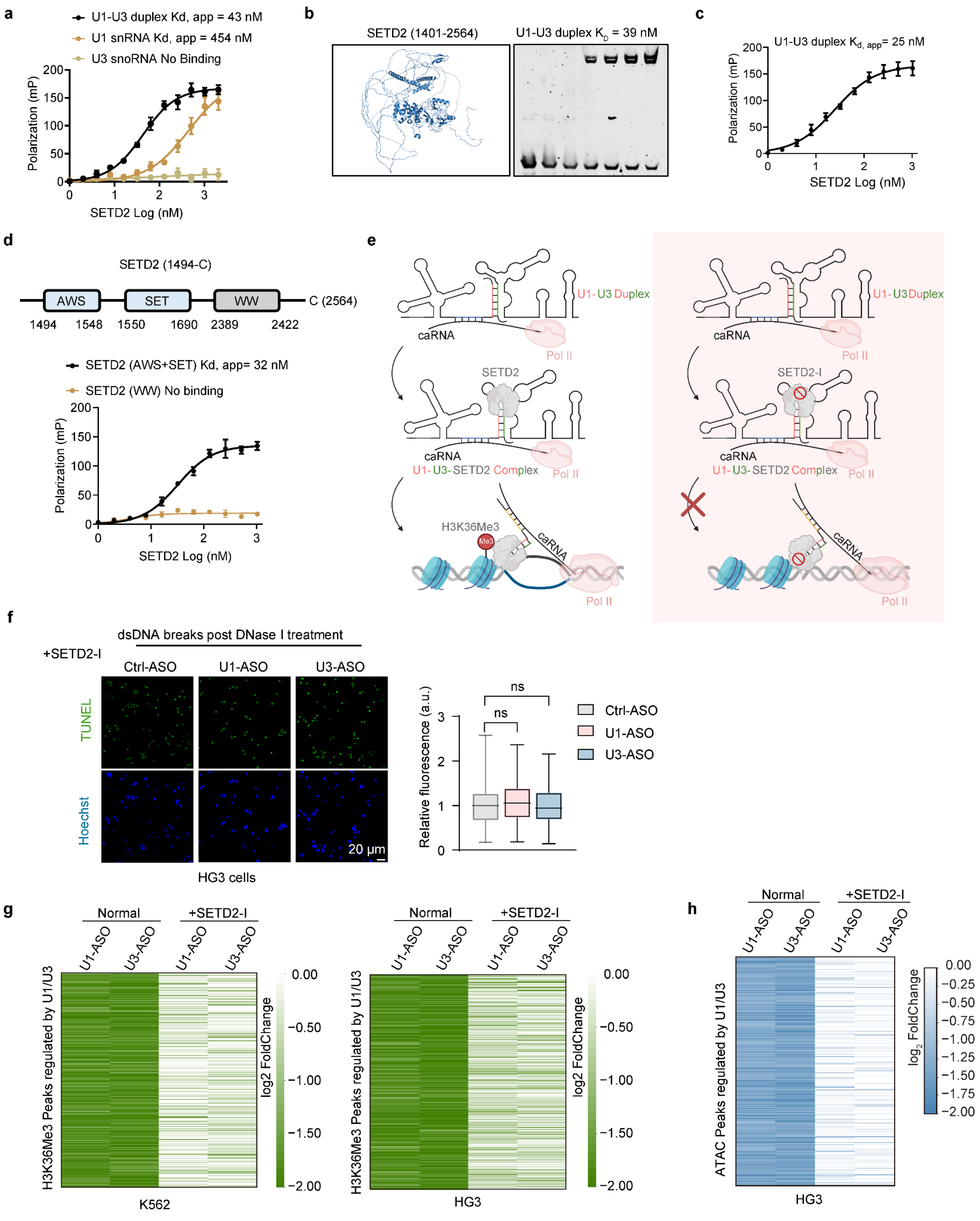


**Extended Data Fig. 7 | SETD2 dependence of U1–U3–mediated chromatin regulation.**
**a**, Fluorescence polarization assay comparing SETD2 binding to U1 snRNA, U3 snoRNA, and the U1–U3 duplex. **b**, Electrophoretic mobility shift assays showing direct binding of recombinant SETD2 truncation mutants to U1 snRNA, U3 snoRNA, and the U1–U3 duplex. **c**,

Fluorescence polarization assay measuring the binding of the U1–U3 RNA duplex to truncated SETD2. **d**, Domain organization of the SETD2 C-terminal region (amino acids 1494–2564), indicating the AWS, SET and WW domains (left), and fluorescence polarization assays measuring binding of the U1–U3 duplex to SETD2 fragments containing AWS+SET or the isolated WW domain (right). **e**, Schematic illustrating that inhibition of SETD2 prevents deposition of H3K36me3 at U1–U3 duplex–recruited sites. **f**, Representative DNase I and TUNEL staining in HG3 cells showing chromatin changes after SETD2 inhibition (left), with quantification across six fields of view (right). **g**, Changes of H3K36me3 signals in K562 and HG3 cells reduced by U1-ASO or U3-ASO before and after SETD2 inhibition. **h**, Changes in ATAC-seq peaks reduced by U1-ASO or U3-ASO before and after SETD2 inhibition in HG3 cells. $n = 3$ biological per group. For **f**, the box plots show the median (centre line), upper and lower quartiles (box limits) and 1–99% (whiskers). Data are mean ± s.d (**a**, **c**, and **d**). *P* value was determined using an unpaired two-tailed *t* test (**f**). TUNEL intensity was quantified by ImageJ.

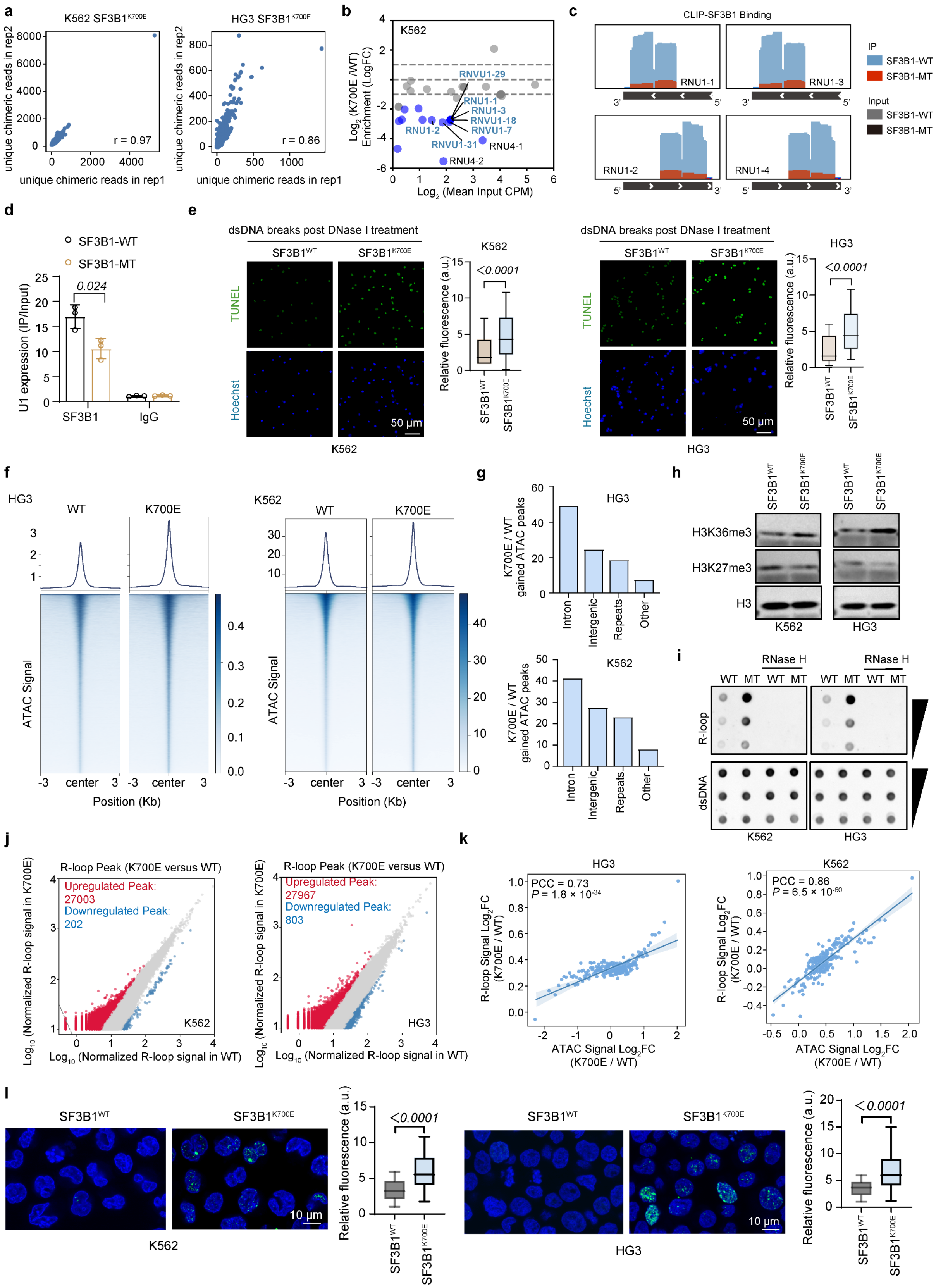
a
K562 SF3B1K700E
HG3 SF3B1K700E
unique chimeric reads in rep2
unique chimeric reads in rep1
r = 0.97
r = 0.86
b
K562
Log2 (K700E /WT) Enrichment (LogFC)
Log2 (Mean Input CPM)
RNVU1-29
RNU1-1
RNU1-3
RNVU1-18
RNVU1-7
RNU1-2
RNVU1-31
RNU4-1
RNU4-2
c
CLIP-SF3B1 Binding
IP
SF3B1-WT
SF3B1-MT
Input
SF3B1-WT
SF3B1-MT
RNU1-1
RNU1-3
RNU1-2
RNU1-4
d
SF3B1-WT
SF3B1-MT
U1 expression (IP/Input)
0.024
SF3B1
IgG
e
dsDNA breaks post DNase I treatment
SF3B1WT
SF3B1K700E
TUNEL
Hoechst
50 μm
K562
HG3
Relative fluorescence (a.u.)
<0.0001
f
HG3
K562
WT
K700E
ATAC Signal
center
Position (Kb)
g
HG3
K562
K700E / WT gained ATAC peaks
Intron
Intergenic
Repeats
Other
h
SF3B1WT
SF3B1K700E
H3K36me3
H3K27me3
H3
K562
HG3
i
RNase H
WT MT WT MT
R-loop
dsDNA
K562
HG3
j
R-loop Peak (K700E versus WT)
Upregulated Peak: 27003
Downregulated Peak: 202
Upregulated Peak: 27967
Downregulated Peak: 803
Log10 (Normalized R-loop signal in K700E)
Log10 (Normalized R-loop signal in WT)
K562
HG3
k
HG3
K562
PCC = 0.73
P = 1.8 × 10-34
PCC = 0.86
P = 6.5 × 10-60
R-loop Signal Log2FC (K700E / WT)
ATAC Signal Log2FC (K700E / WT)
l
SF3B1WT
SF3B1K700E
10 μm
K562
HG3
Relative fluorescence (a.u.)
<0.0001

**Extended Data Fig. 8 | Chromatin alterations and genome instability in SF3B1-mutant cells.**

**a**, Correlation of unique chimeric reads obtained from biological replicates across HG3 $SF3B1^{K700E}$ and K562 $SF3B1^{K700E}$ cell lines. **b**, CLIP–seq analysis showing differential snRNA binding in K562 $SF3B1^{K700E}$ (MT) versus K562 $SF3B1^{K700K}$ (WT) cells. Scatter plot displays $\log_2$ enrichment changes (MT/WT) relative to mean input abundance for individual snRNAs. Blue points indicate snRNAs with reduced association in mutant cells. **c**, CLIP-seq using anti-Flag antibody showing RNA binding profiles of K562 $SF3B1^{K700K}$ and $SF3B1^{K700E}$ cells, respectively, with IGV tracks depicting U1-binding sites. **d**, RIP-qPCR showing enrichment of U1 snRNA by Flag antibody in K562 $SF3B1^{WT}$ and $SF3B1^{K700E}$ cells. **e**, Representative DNase I and TUNEL staining in $SF3B1^{WT}$ and $SF3B1^{K700E}$ K562 cells as well as HG3 cells (left), with quantification across six fields of view (right). **f**, Average ATAC-seq profiles and heat maps of chromatin accessibility in $SF3B1^{WT}$ and $SF3B1^{K700E}$ K562 cells as well as HG3 cells. **g**, Genomic distribution of ATAC-seq peaks gained in HG3 and K562 $SF3B1^{K700E}$ cells versus HG3 and K562 $SF3B1^{WT}$ cells. **h**, Western blot showing H3K36me3 and H3K27me3 differences between $SF3B1^{WT}$ and $SF3B1^{K700E}$ K562 and HG3 cells. **i**, Dot blot showing R-loop levels in $SF3B1^{WT}$ and $SF3B1^{K700E}$ K562 and HG3 cells. **j**, Correlation between differential R-loop peaks and ATAC-seq peaks in $SF3B1^{WT}$ and $SF3B1^{K700E}$ K562 and HG3 cells. **k**, Correlation between changes in chromatin accessibility and R-loop abundance in SF3B1-K700E K562 and HG3 cells. Scatter plots show the relationship between ATAC-seq $\log_2$ fold change and R-loop signal $\log_2$ fold change in HG3 (left) and K562 (right) cells. **l,** Representative images and quantification of phosphorylated γ-H2AX foci in $SF3B1^{WT}$ and $SF3B1^{K700E}$ K562 and HG3 cells. Quantification was performed across six fields of view. $n = 3$ biological per group. Data are mean ± s.d for **d**. $P$ value was determined using an unpaired two-tailed $t$ test (**d**, **e**), and Pearson's correlation coefficient (PCC) was used to assess correlation (**k**). Signal intensity was measured with ImageJ.

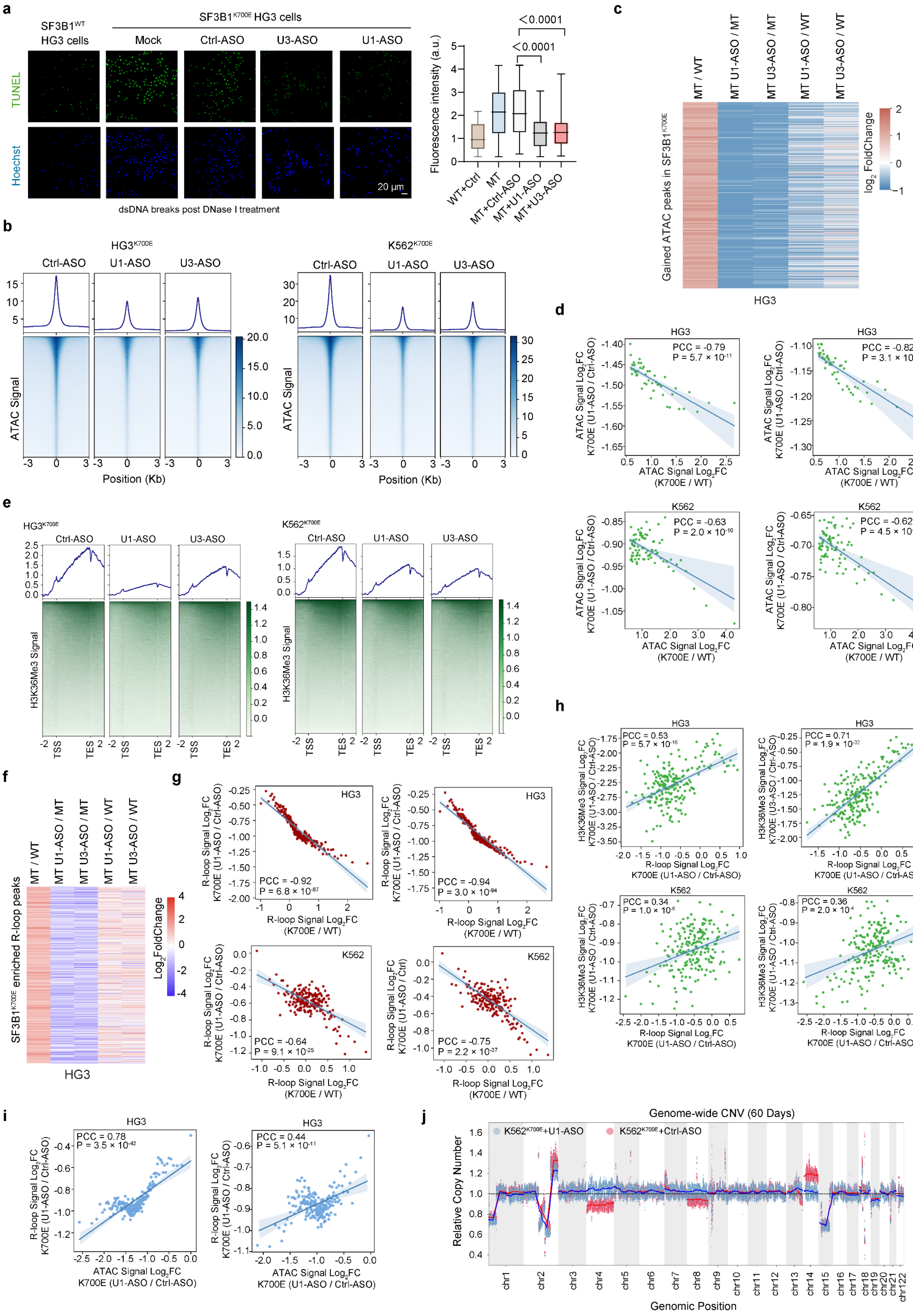
a
SF3B1WT HG3 cells
SF3B1K700E HG3 cells
Mock
Ctrl-ASO
U3-ASO
U1-ASO
TUNEL
Hoechst
20 μm
dsDNA breaks post DNase I treatment
Fluorescence intensity (a.u.)
<0.0001
<0.0001
WT+Ctrl
MT
MT+Ctrl-ASO
MT+U1-ASO
MT+U3-ASO
b
HG3K700E
K562K700E
Ctrl-ASO
U1-ASO
U3-ASO
ATAC Signal
Position (Kb)
c
MT / WT
MT U1-ASO / MT
MT U3-ASO / MT
MT U1-ASO / WT
MT U3-ASO / WT
Gained ATAC peaks in SF3B1K700E
log2 FoldChange
HG3
d
HG3
K562
PCC = -0.79
P = 5.7 × 10-11
PCC = -0.82
P = 3.1 × 10-12
PCC = -0.63
P = 2.0 × 10-10
PCC = -0.62
P = 4.5 × 10-10
ATAC Signal Log2FC K700E (U1-ASO / Ctrl-ASO)
ATAC Signal Log2FC (K700E / WT)
e
HG3K700E
K562K700E
H3K36Me3 Signal
TSS
TES
f
SF3B1K700E enriched R-loop peaks
Log2FoldChange
HG3
g
HG3
K562
PCC = -0.92
P = 6.8 × 10-87
PCC = -0.94
P = 3.0 × 10-94
PCC = -0.64
P = 9.1 × 10-25
PCC = -0.75
P = 2.2 × 10-37
R-loop Signal Log2FC K700E (U1-ASO / Ctrl-ASO)
R-loop Signal Log2FC K700E (U1-ASO / Ctrl)
R-loop Signal Log2FC (K700E / WT)
h
HG3
K562
PCC = 0.53
P = 5.7 × 10-16
PCC = 0.71
P = 1.9 × 10-32
PCC = 0.34
P = 1.0 × 10-6
PCC = 0.36
P = 2.0 × 10-4
H3K36Me3 Signal Log2FC K700E (U1-ASO / Ctrl-ASO)
H3K36Me3 Signal Log2FC K700E (U3-ASO / Ctrl-ASO)
R-loop Signal Log2FC K700E (U1-ASO / Ctrl-ASO)
i
HG3
PCC = 0.78
P = 3.5 × 10-42
PCC = 0.44
P = 5.1 × 10-11
R-loop Signal Log2FC K700E (U1-ASO / Ctrl-ASO)
ATAC Signal Log2FC K700E (U1-ASO / Ctrl-ASO)
j
Genome-wide CNV (60 Days)
K562K700E+U1-ASO
K562K700E+Ctrl-ASO
Relative Copy Number
Genomic Position

**Extended Data Fig. 9 | U1–U3 disruption modulates chromatin accessibility, R-loop accumulation and genome instability in SF3B1$^{K700E}$ cells.**

**a,** Representative DNase I and TUNEL staining in HG3 SF3B1$^{K700E}$ cells after U1-ASO or U3-ASO treatment (left), with quantification across six fields of view (right). TUNEL intensity was quantified by ImageJ. **b,** Average ATAC-seq profiles and heat maps of chromatin accessibility in K562 SF3B1$^{K700E}$ and HG3 SF3B1$^{K700E}$ cells following U1-ASO or U3-ASO treatment. **c,** Heat map illustrating the spike-in-calibrated ATAC–seq signals across ATAC-seq peak regions that are more accessible in HG3 SF3B1$^{K700E}$ cells than in HG3 SF3B1$^{WT}$ cells, and these signal changes in HG3 SF3B1$^{K700E}$ cells following U1-ASO or U3-ASO treatment. **d,** Correlation analysis comparing ATAC-seq changes upon U1–U3 disruption (U1-ASO/Ctrl-ASO) with ATAC-seq alterations observed in SF3B1$^{K700E}$ versus SF3B1$^{WT}$ K562 and HG3 cells. **e,** CUT&Tag profiles of H3K36me3 following U1-ASO or U3-ASO treatment in SF3B1$^{K700E}$ K562 and HG3 cells. **f,** Heatmap showing spike-in–calibrated R-loop signals at R-loop peak regions that are increased in HG3 SF3B1$^{K700E}$ cells compared with HG3 SF3B1$^{WT}$cells, and their changes following U1-ASO or U3-ASO treatment. **g,** Correlation analysis comparing R-loop signal changes upon U1–U3 disruption (U1-ASO/Ctrl-ASO) with R-loop alterations observed in SF3B1$^{K700E}$ versus SF3B1$^{WT}$ K562 and HG3 cells. **h,** Correlation analysis comparing H3K36me3 changes induced by U1–U3 disruption (U1-ASO/Ctrl-ASO) with corresponding R-loop alterations in SF3B1$^{K700E}$ versus SF3B1$^{WT}$ K562 and HG3 cells. **i,** Correlation between peaks showing reduced ATAC-seq signal and reduced R-loop accumulation after ASO treatment in HG3 SF3B1$^{K700E}$ cells. **j,** Whole-genome sequencing (WGS) analysis of copy-number variation (CNV) in K562 SF3B1$^{K700E}$ cells 60 days after U1-ASO or Ctrl-ASO treatment. CNV profiles were derived from WGS read-depth data and normalized to Ctrl-ASO–treated K562 SF3B1$^{K700K}$ data. For **a**, the box plots show the median (centre line), upper and lower quartiles (box limits) and 1–99% (whiskers). Pearson's correlation coefficient (PCC) was used to assess correlation (**d**, **g**, **h** and **i**). *P* value was determined using an unpaired two-tailed *t* test (**a**). TUNEL intensity was quantified by ImageJ.

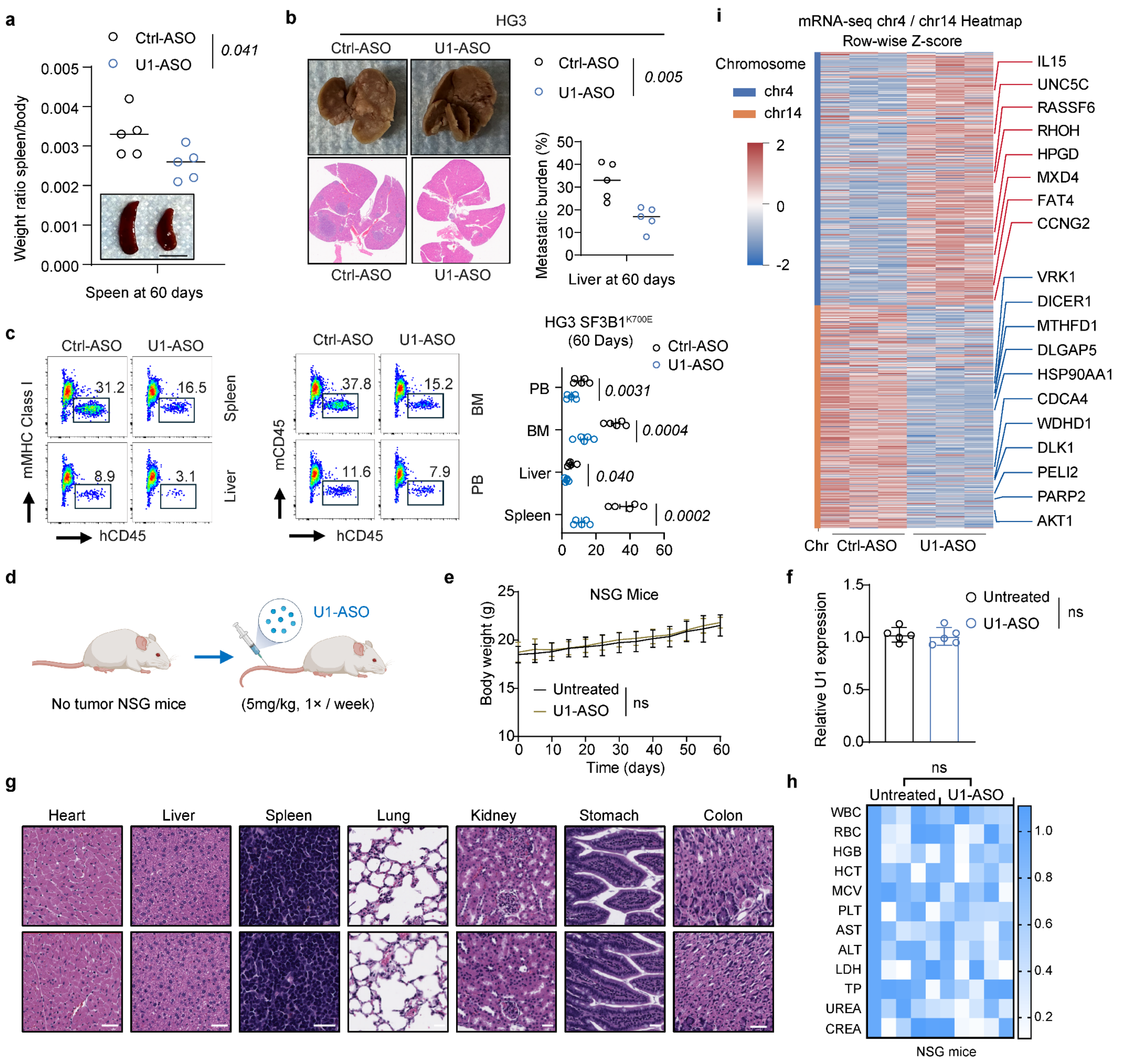

**Extended Data Fig. 10 | In vivo effects and safety of U1-ASO treatment.**
**a**, Representative images showing spleen size in control-ASO– and U1-ASO–treated mice bearing HG3 SF3B1$^{K700E}$ xenografts. Quantification of spleen weight normalized to body weight is shown on the top. **b**, Number of hepatic metastatic foci in control-ASO– and U1-ASO–treated mice bearing HG3 SF3B1$^{K700E}$ xenografts. **c**, Representative flow cytometry plots showing human leukemia cell (HG3 SF3B1$^{K700E}$) infiltration in peripheral blood, bone marrow, and solid organs. **d**, Schematic of the experimental design for evaluating U1-ASO safety in mice. **e**, Body weight trajectories of NSG mice treated with U1-ASO or left untreated over the indicated time course. **f**, Relative U1 snRNA expression in mouse peripheral blood measured by RT–qPCR in untreated and U1-ASO–treated mice. **g**, Representative haematoxylin and eosin (H&E) staining of major organs (heart, liver, spleen, lung, kidney, stomach and colon; scale bar, 50 μm). **h**, Safety analysis of untreated and U1-ASO–treated mice, showing blood biochemistry and liver and kidney function. **i**, Heat map of differential mRNA expression from chromosomes 4 and 14

in xenograft-derived K562 $SF3B1^{K700E}$ cells, shown as row-wise z-scores. Representative upregulated and downregulated genes are highlighted with red and blue labels, respectively. $n$ = 5 biological per group. Data are mean ± s.d. $P$ value was determined using an unpaired two-tailed $t$ test (**a**, **b**, **c**, **f** and **h**) or two-way ANOVA with Tukey's post-test (**e**). ns indicates no statistically significant difference.
